# Evolution-Inspired Engineering of Anthracycline Methyltransferases

**DOI:** 10.1101/2021.08.05.455194

**Authors:** Pedro Dinis, Heli Tirkkonen, Benjamin Nji Wandi, Vilja Siitonen, Jarmo Niemi, Thadée Grocholski, Mikko Metsä-Ketelä

## Abstract

*Streptomyces* soil bacteria produce hundreds of anthracycline anticancer agents with a relatively conserved set of genes. This diversity depends on the rapid evolution of biosynthetic enzymes to acquire novel functionalities. Previous work has identified *S*-adenosyl-L-methionine-dependent methyltransferase-like proteins that catalyze either 4-O-methylation, 10-decarboxylation or 10-hydroxylation, with additional differences in substrate specificities. Here we focused on four protein regions to generate chimeric enzymes using sequences from four distinct subfamilies to elucidate their influence in catalysis. Combined with structural studies we managed to depict factors that influence gain-of-hydroxylation, loss-of-methylation and substrate selection. The engineering expanded the catalytic repertoire to include novel 9,10-elimination activity, and 4-O-methylation and 10-decarboxylation of unnatural substrates. The work provides an instructive account on how the rise of diversity of microbial natural products may occur through subtle changes in biosynthetic enzymes.

## Introduction

Microbial natural products are found ubiquitously from the biosphere, where they are present in enormous numbers and display tremendous diversity in their chemical structures. In particular, *Streptomyces* soil bacteria have tens of thousands of secondary metabolic pathways for production of natural products^1^. The potent bioactivities have made microbial natural products a cornerstone of modern medicine ever since the discoveries of penicillin and streptomycin^2^. Approximately two thirds of antibiotics and one third of anticancer agents are natural products or their semi-synthetic derivatives^3^. The full scope of microbial chemodiversity is still unknown, as the producing organisms have been noted to harbour an excess of biosynthetic gene clusters in relation to the number of metabolites observed under monocultures^4^. Many secondary metabolic pathways have been thought to be silent and require specific environmental cues for activation^5^. Microbial natural products typically function as chemical warfare agents against competing organisms^6^, but a fraction are involved in more complex chemical communication interactions^7^. Both intra- and inter-species chemical communication, which may be either agonistic or antagonistic in nature, has been reported^6,8^. Even trans-kingdom interactions have been demonstrated, such as production of the terpenoid geosmin by *Streptomyces* to attract arthropods to assist is spore dispersal^9^.

From the perspective of evolutionary biochemistry, the most remarkable aspect of microbial natural products is the divergent enzyme evolution responsible for the emergence of chemodiversity. Several models for the molecular evolution of secondary metabolism have been presented^10-12^, which highlight important differences to the evolution of the conserved central or primary metabolic pathways. A key concept dictates that the appearance of novel bioactive molecules provides advantages for the producing organisms under shifting environmental conditions and selective pressures^8,13^. Proteins involved in secondary metabolism may be considered generalists^14,15^, which manifests as promiscuity^16^ and slow reactions rates^17^. The situation is in contradiction to central metabolism, where proteins catalyse their reactions both efficiently and specifically to meet the demands for high metabolic fluxes^10,18^. Furthermore, purifying selection has shaped central metabolism by improving the catalytic properties of ancestral generalist enzymes at the cost of reducing genetic variation^11,19^. However, resilience towards purifying selection has allowed enzymes involved in secondary metabolism to remain in a generalist state and acquire atypical catalytic abilities above and beyond what has been classically thought possible.

An example of microbial chemical diversity are the anthracyclines, which are glycosylated tetracyclic aromatic polyketides^20^ (Fig. 1a). Anthracyclines are mainly produced by Actinobacteria and to date 408 metabolites have been described^21^. The complexity of anthracyclines is generated in the so called tailoring steps of the biosynthesis, where the carbon framework is modified in many ways, and in glycosylation steps that differ in the type and number of carbohydrate units attached to the aglycones^22^. The importance of anthracyclines is in their potent cytotoxic activity against tumour cells, and metabolites such as aclacinomycin and doxorubicin are widely used in cancer chemotherapy^23^. Although the biological activity is complex, poisoning of topoisomerases and intercalation to DNA are important contributors^24,25^. However, other factors such as formation of reactive oxygen species^26^, the ability to evict histones from chromatin^27^ and proteolytic activation of transcription factors^28^ have been noted.

**Figure 1.**
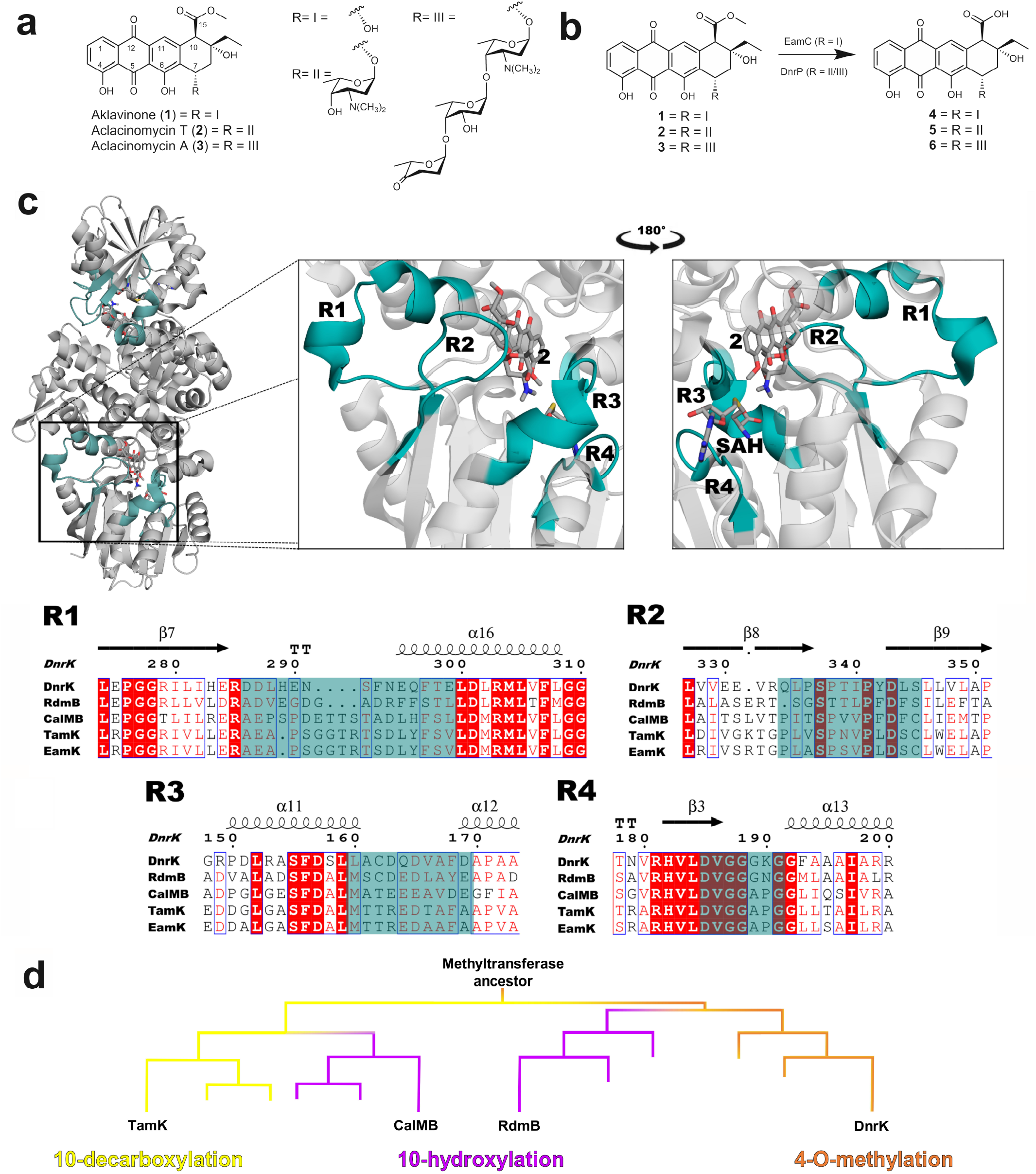
Diversity of SAM-dependent methyltransferase-like proteins in anthracycline biosynthesis. **a,** Structures of aklavinone (**1**), aclacinomycin T (**2**) and aclacinomycin A (**3**). **b,** An initial 15-methylesterase activity is required for the modification reactions at C-10 position under study. EamC was used with **1**, while DnrP was used with both **2** and **3**. **c,** Structure of DnrK, with exchanged regions (R1-R4) highlighted (teal). Multiple sequence alignment (ClustalOmega^49^, ENDScript^50^) of the relevant members of the family, with chimeric regions highlighted (teal), **d,** Evolutionary tree of the proteins under study. 4-O-methyltransferase DnrK clade (orange) shows the ancestral 4-*O*-methylation activity of monoglycosylated substrates (**5**), as well as moonlighting 10-decarboxylation activity. 10-decarboxylase TamK (yellow) has lost the ancestral methylase activity and evolved to accept **4** and **5** as substrates. Convergent evolution has led to gain of 10-hydroxylation activity (magenta), together with broadening of substrate specificity to accommodate **4, 5** and **6** as substrates.

Tailoring steps catalysed by SAM methyltransferase-like proteins^29^ have yielded particular insight into evolution of anthracyclines and generation of structural diversity. DnrK is a canonical *S*-adenosyl-L-methionine (SAM) -dependent methyltransferase that catalyses 4-O-methylation (Fig. 2a) in daunorubicin biosynthesis^30^. Recent studies have uncovered that DnrK is bifunctional and catalyses additional 10-decarboxylation as a moonlighting activity^31^. However, the activity depends on the presence of a free 10-carboxyl group in the substrate generated by aclacinomycin 15-methylesterases such as EamC and DnrP (Fig. 1b)^29^. DnrK is quite specific with respect to the length of the carbohydrate chain at C-7, accepting only monoglycosides (Fig. 1a).

**Figure 2.**
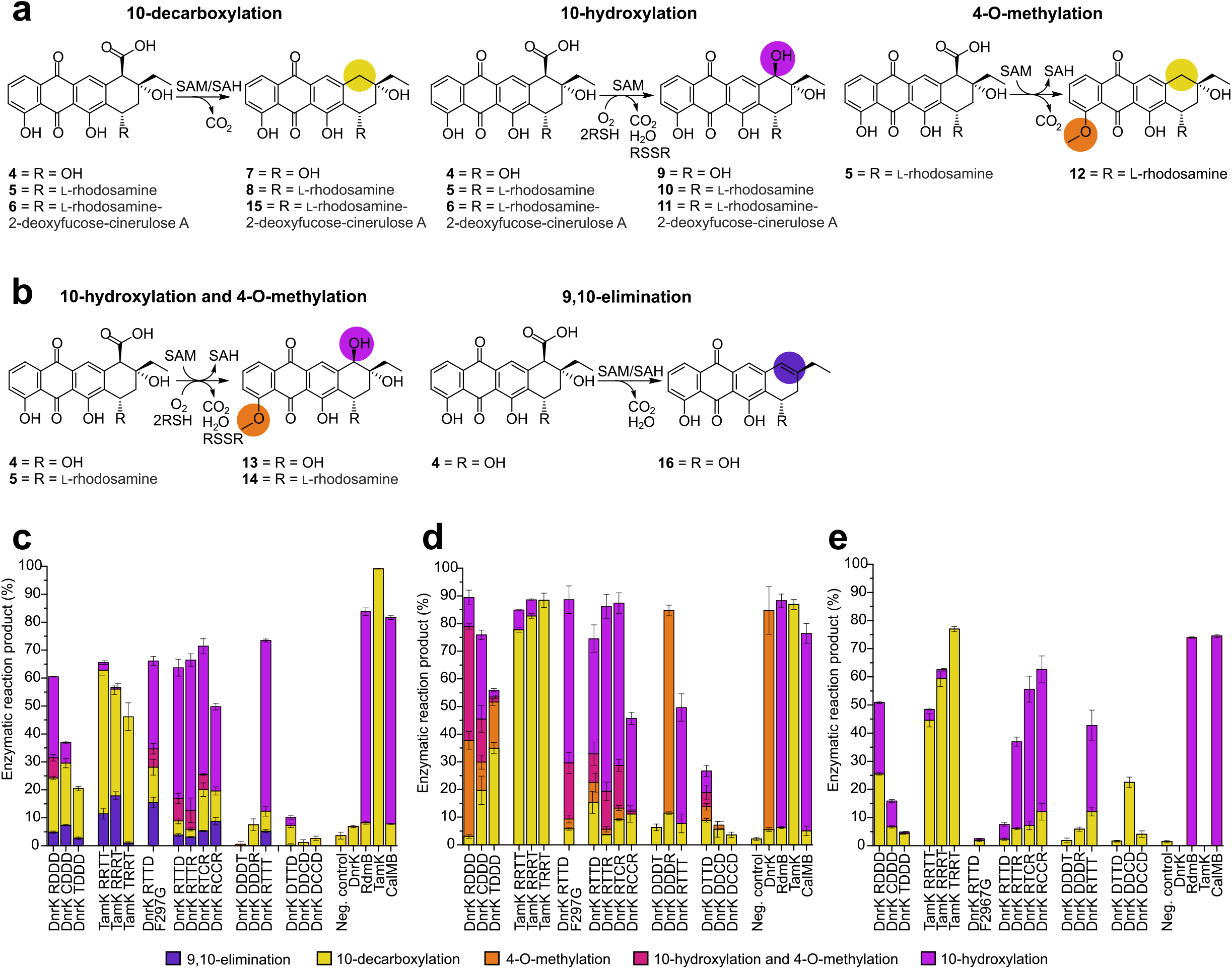
Enzymatic activity measurements of SAM-dependent methyltransferase-like proteins. **a,** Native reactions performed by native or chimeric enzymes, **b,** Non-natural reactions performed by the chimeric enzymes. **c,** Enzymatic reaction products with **4** as a substrate. **d,** Enzymatic reaction products with **5** as a substrate. **e,** Enzymatic reaction products with **6** as a substrate. The reaction product yields are shown as mean value ± standard deviation and were calculated from HPLC chromatogram traces by normalized peaks areas. The overall percentages may not add up to 100 % in all samples meaning that some unreacted substrate remained in the reaction mixture.

Conversely, RdmB (53.4% sequence identity to DnrK) from the ε-rhodomycin pathway is an anthracycline 10-hydroxylase requiring SAM, molecular oxygen and a thiol reducing agent for activity (Fig. 2a)^32^. RdmB harbors broad substrate specificity and is able to turn over aglycones, mono- and triglycosylated anthracyclines^29^ (Fig. 1a). The 10-hydroxylation activity has evolved twice independently, as indicated by the discovery of a second clade of 10-hydroxylating enzymes, such as CalMB (50.8% sequence identity to DnrK), that have a similar tolerance towards glycosylated substrates as RdmB^29^. Finally, the enzyme family was expanded by identification of EamK (52.4% sequence identity to DnrK) from the komodoquinone B pathway, which catalyses solely 10-decarboxylation (Fig. 2a)^29^. The specificity of EamK and related 10-decarboxylase TamK lies between DnrK and RdmB, since anthracycline aglycones and monoglycosylated compounds are accepted as substrates (Fig. 1a).

Here we have taken advantage of the natural diversity of anthracycline methyltransferases and employed extensive chimeragenesis^33^ to probe factors that govern catalysis and substrate recognition. We have mimicked evolutionary processes and generated 24 chimeric proteins using two proteins scaffolds and sequences from four proteins. The chimeras expanded the catalytic repertoire to include 9,10-elimination activity, and broadened the substrate tolerance for 4-O-methylation and 10-decarboxylation. Functional and structural characterization revealed a complex network of interactions that determine the molecular basis for the altered enzymatic activities. The engineering efforts demonstrated how minor changes in generalist proteins may induce drastic effects in their functions as biocatalysts.

### Selection of regions for chimeragenesis

Three regions have previously been noted as critical for the functional diversification of the 10-hydroxylase RdmB from the canonical 4-O-methyltransferase DnrK^31^. The first region R1 (residues 286-299 in DnrK) consists of the N-terminal half of α16 and the preceding loop region. Helix α16 is situated parallel to the tetracyclic anthracycline substrate with R1 residing near the 10-carboxyl group (Fig. 1c). The switch in activity from 4-O-methylation to 10-hydroxylation could be attributed to insertion of a single S298 residue in DnrK into the α16 helix, which triggered rotation of the neighbouring F297 towards the ligand^31^. This shift resulted in a solvent-free active site architecture reminiscent of RdmB, where a channel leading to the surface of the protein is closed by F300. A mechanistic model for 10-hydroxylation has proposed that exclusion of water molecules from the active site allows an anthracycline 10-carbanion intermediate to react with molecular oxygen for 10-hydroxylation to occur^31^. In turn, region R2 (residues 334-345) comprises the loop region between β8 and β9 that folds over the active site and is in contact with the ligand (Fig. 1c). Finally, the loop region between α11 and α12 (residues 160-169), which interacts with the carbohydrate unit of the substrate, was defined as region R3 (Fig. 1c). Regions R2 and R3 from RdmB were found to expand the substrate tolerance of DnrK towards triglycosylated anthracyclines^31^. The regions R1-R3 described here are highly variable in the four clades of anthracycline methyltransferases-like proteins (Fig. 1c) and hence remained unchanged in this study.

The expansion of the family by the 10-decarboxylases directed us to differences in the loop region between β3 and α13, which is located close to the binding site of the co-substrate SAM. The 4-O-methylation of DnrK has strict requirements for correct geometrical positioning of the substrate and SAM for an S_N_2 reaction to occur^34^ and misalignment of the ligands has been proposed to be the reason why RdmB does not have methylation activity^32^. Since this may be triggered either by shifting the position of the substrate or the co-substrate, we selected the SAM binding site as region R4. Noteworthy, the 10-decarboxylases EamK and TamK harbor two point mutations G189A/K190P in R4 in comparison to DnrK, which is interesting as the presence of a proline restricts backbone conformations^35^.

Due to the complex nature of the constructs, we devised a simple nomenclature to describe the chimeras. In this system, the scaffold enzyme is firstly mentioned, followed by a sequence of four letters, corresponding to the origin of the regions R1-R4 (D for DnrK, R for RdmB, C for CalMB and T for TamK). To exemplify, TamK DRRD is a chimera with a TamK scaffold, regions R1 and R4 from DnrK and regions R2 and R3 from RdmB.

### Enzymatic activities of the chimeric proteins

The great number of combinatorial possibilities with four regions (R1-R4) and four possible donor sequences (DnrK, RdmB, CalMB and TamK) precluded a systematic survey. The chimeras selected for the study were designed based on structural information to probe research questions with 12 chimeras created using DnrK as the main scaffold. In addition, the viability of 10-decarboxylases to serve as scaffolds was tested with eight unique chimeras using TamK (Supplementary Table 1). No hydroxylases were selected due to previous challenges in engineering the enzymatic activity of RdmB^31^.

The relative enzymatic activities were tested with three compounds: the aglycone aklavinone (**1**); the monoglycosidic aclacinomycin T (**2**); and the triglycosidic aclacinomycin A (**3**) (Fig. 1a). In order to allow 10-decarboxylation and 10-hydroxylation to occur, all reactions were preceded by a 15-methylesterase reaction either by EamC^29^ or DnrP^36^. Subsequently intermediates **4, 5** and **6** were used as substrates for the chimeras together with the reducing agent dithiotreitol (DTT) and SAM.

Analysis of reaction products by HPLC revealed that 16 chimeras harboured diverse enzymatic activities (Fig. 2c-2e). The majority of reaction products could be identified by comparison to standards obtained from previous studies, which included 10-decarboxylation (**7, 8, 15**), 10-hydroxylation (**9, 10, 11**) and 4-O-methylation (**12**) products, and compounds with both 4-O-methyl and 10-hydroxyl groups (**13, 14**) (Supplementary Figs. 1-7). To confirm the identity of **15**, which was produced by several TamK-based chimeras when **6** was used as a substrate, the glycosidic units were acid hydrolyzed and the resulting aglycone **7** was compared to an authentic standard 7 (Supplementary Fig. 8). MS measurements supported that **15** was a triglycosylated 10-decarboxylation reaction product ([M+H]^+^, ESI+ obs. 754.3481, calc. 754.3433, Supplementary Fig. 9). Similarly, four chimeras appeared to expand the substrate specificity for 4-O-methylation and 10-hydroxylation towards **4** based on MS analysis ([2M+Na]^+^, ESI+ obs. 791.2254, calc. 791.2310, Supplementary Fig. 10). In order to identify the novel product **13**, we removed the L-rhodosamine aminosugar from standard **14** by acid hydrolysis and compared the products by LC-MS-EICS (Supplementary Fig. 11).

We also noted that ten chimeras produced a novel product **16**, which appeared to be an elimination product based on HR-MS measurement ([M-H]^-^, ESI-obs. 335.0923, calc. 335.0925, Supplementary Fig. 12), when **4** was used as a substrate. The highest yields were detected with TamK RRRT, which was used in large-scale reactions to obtain sufficient material for structure elucidation. NMR-experiments (^1^H, ^13^C, HSQCDE, COSY and HMBC) revealed that an additional 9,10-elimination reaction had occurred in **16** (Supplementary Table 2 and Supplementary Figs. 13-18).

### The critical role of region R1 in reaction type determination

Previous studies have demonstrated the importance of R1 from RdmB in gain-of-hydroxylation activity of DnrK RDDD with **5** and **6**^31^. At the time, the activity could not be assayed for **4**, since the 15-methylesterase DnrP does not accept **1** as a substrate. However, the recent discovery of the 15-methylesterase EamC^29^ allowed us to show that DnrK RDDD is able accept **4** with 29 % hydroxylation reactivity (Fig. 2c).

Here we wished to focus on the second clade of 10-hydroxylating enzymes, since R1 of CalMB differs greatly from both DnrK and RdmB (Fig. 1c). Region R1 in CalMB is extended by an insertion of three residues and has a Phe to His substitution at the critical site in comparison to RdmB. Despite these differences, DnrK CDDD behaved similar to its R1 provider with the highest yield of hydroxylation product (46 %) shown with **5** (Fig. 2d).

Next, we introduced R1 from the 10-decarboxylase TamK, creating DnrK TDDD. The region is reminiscent to CalMB in terms of length, but contains a Tyr residue instead of His at the key junction. Activity assays confirmed the importance of R1 in determination of reaction chemistry, since the 10-decarboxylation activity of DnrK TDDD increased with all substrates, particularly with **5** (35 %), in comparison to DnrK RDDD and DnrK CDDD (Fig. 2d).

The use of TamK as a protein scaffold proved to be moderately successful in highlighting the role of R1 in catalysis. TamK RRTT and TamK RRRT acquired minor 10-hydroxylation activity as expected based on the R1 donor sequence, while TamK TRRT showed no 10-hydroxylation profile (Fig. 2c-2e).

Similarly to experiments with the RdmB scaffold^31^, engineering gain-of-methylation functionality was challenging, as chimeras TamK DRRD, TamK DRTD, TamK DDRD, TamK DDTD and TamK DDDD did not display any measurable enzymatic activity.

### Structural basis for 10-decarboxylation activity

All proteins from the four phylogenetic clades share 10-decarboxylation as a common feature. In addition to the true 10-decarboxylase TamK^29^ and the moonlighting 10-decarboxylase activity of DnrK^31^, formation of a 10-carbanion intermediate has been proposed to be a key step in the 10-hydroxylation activity of RdmB^32^. A fully conserved arginine R303 (DnrK numbering) residue (Fig. 1c), which is within hydrogen bonding distance from the 10-carboxyl group, is considered to initiate the reaction^31,32^.

Structural determination of DnrK TDDD revealed that insertion of three additional residues in R1 extends the α16 helix by one additional half-turn in comparison to DnrK and DnrK S298 (Fig. 3a-3c). However, the longer helix protrudes away from the active site towards bulk solvent and is unlikely to have a significant influence in catalysis. DnrK contains Q296 that allows access for solvent ions to reach the active site through an open channel (Fig. 3a), which is blocked by F297 in DnrK S298 (Fig. 3b). DnrK TDDD harbours Y299 in the structurally equivalent position (Fig. 3c), but the solvent channel remains open due to an additional interaction of Y299 with the ligand; a hydrogen bond between the tyrosine and the carbonyl oxygen of the substrate rotates the side chain of Y299 in a manner that prevents channel closure (Fig. 3c).

**Figure 3.**
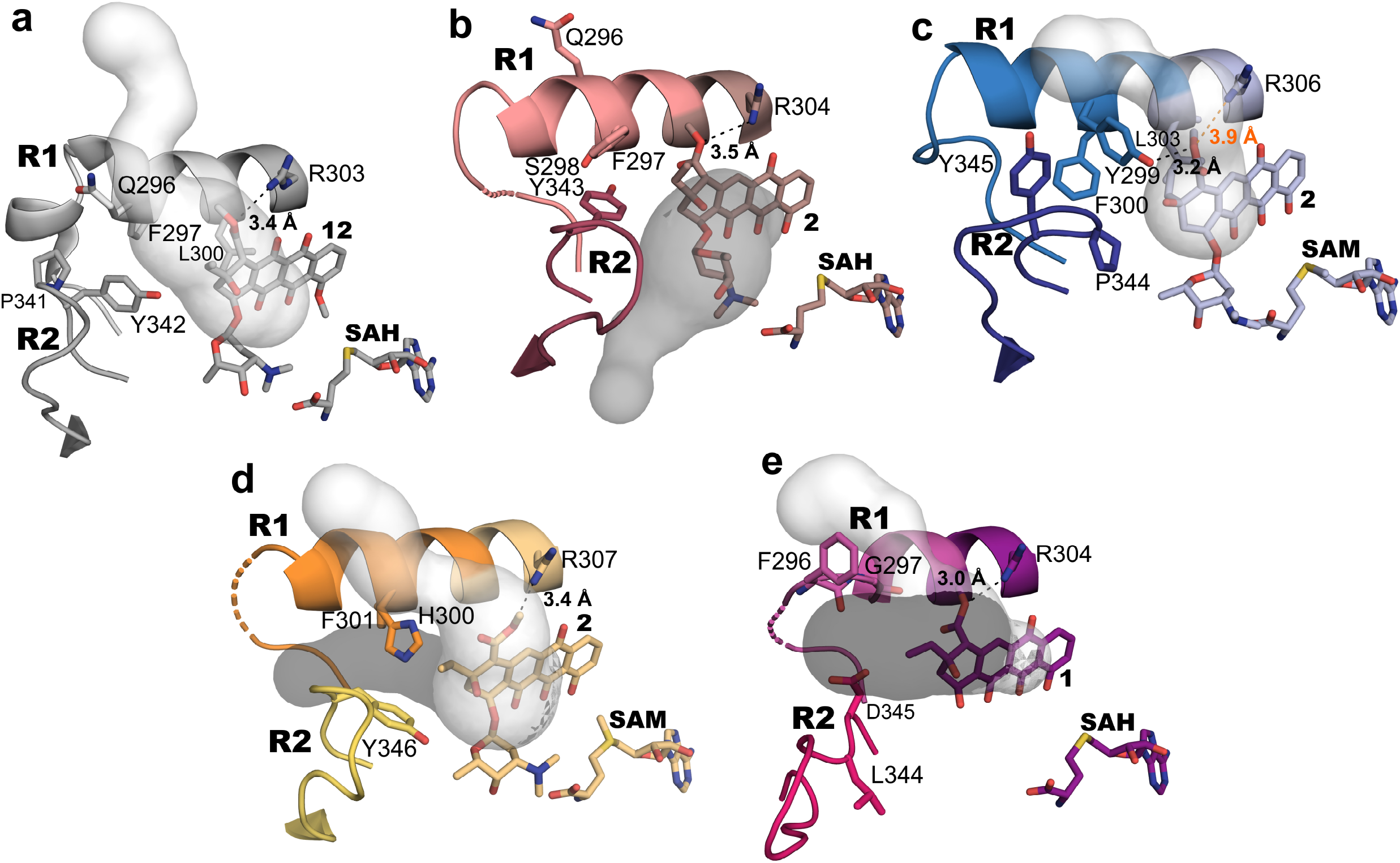
Structural analysis of regions R1 and R2 from different clades. Region R1 displays noticeable differences, particularly in regards to positioning of aromatic amino acids that are involved in formation of a water channel towards bulk solvent (light grey). In addition, movement of R2 depicts closed and open conformations of the chimeras, where a large channel is formed (dark grey) to provide access for the substrate to the active site. **a,** DnrK (PDBID: 1TW2) contains an open solvent channel due to Q296 that prevents 10-hydroxylation. **b,** DnrK S298 (PDBID: 4WXH) shows the closure of the channel by F297 to prevent access of bulk solvent to the active site. **c,** The solvent channel remains open in DnrK TDDD due to interaction of Y299 with the ligand **2**. **d,** DnrK CDDD crystallized in an open conformation, with a substrate access channel (dark grey) visible between regions R1 and R2. Subtle movement of H300 and Y346 is required to close the solvent channel (light grey) in the closed conformation. **e,** The dramatic rearrangements in DnrK RTTD F297G lead to repositioning of a neighbouring residue F296 towards the solvent channel (light grey). This structure is in an open conformation with a visible substrate access channel (dark grey).

The residue Y299 is fully conserved in 10-decarboxylating enzymes and it may additionally have a mechanistic role in activity. Similarly to a methylmalonyl-CoA decarboxylase^37^, Y299 forms a polar contact with the carboxylate unit for the correct positioning of the leaving group. L303 and P344, which are conserved in all members of this family (L300 and P341 DnrK Numbering, Figure 3a), are in DnrK TDDD responsible for a hydrophobic environment that destabilizes the ground state of the substrate, favouring the release of neutral carbon dioxide molecule^37,38^.

The residue Y299 is fully conserved in 10-decarboxylating enzymes and it may additionally have a mechanistic role in activity. Similarly to a methylmalonyl-CoA decarboxylase^37^, Y299 forms a polar contact with the carboxylate unit for the correct positioning of the leaving grouphich are conserved in all members of this family, are responsible for a hydrophobic environment that destabilizes the ground state of the substrate, favouring the release of neutral carbon dioxide molecule^37,38^.

### Structural basis for 10-hydroxylation activity

To investigate if the previously proposed mechanism for 10-hydroxylation^31^ holds true for the second clade of 10-hydroxylating enzymes, we solved the structure of DnrK CDDD. The structure demonstrates that two aromatic residues, H300 and Y346, are positioned adjacent to the channel. The analysis is complicated by the fact that DnrK CDDD crystallized in an open conformation where R2 hosting Y346 has moved away from R1 and, as a consequence, H300 does not fully close the solvent channel (Fig. 3d). However, it seems likely that movement of R2 to close the active site will lead to rearrangement of the H300:Y346 interaction. A similar effect can be observed in the reported structures of DnrK, where significant differences in R2 emerged between open and closed conformations^39^.

To overcome the experimental uncertainty, we replaced the critical phenylalanine residue with glycine in DnrK RTTD F297G. According to our mechanistic model, removal of the bulky hydrophobic side chain would allow bulk solvent to access the active site and prevent 10-hydroxylation^31^. Unexpectedly the mutant displayed moderate 10-hydroxylation activity towards **4** and **5** with 38 and 79 % conversion, respectively, with limited reactivity towards **6** (Fig. 2c-2e). The contradiction was cleared by structural data from DnrK RTTD F297G, which revealed significant rearrangements in the active site architecture. The introduction of G297 had induced a disruption to the α16 helix secondary structure and, consequently, lead to rotation of a different phenylalanine residue (the preceding F296) towards the active site. While cavity analysis shows an open conformation also for this structure, F296 is ideally positioned to block the solvent channel in the closed conformation (Fig. 3e). The data therefore provides independent support for the importance of solvent exclusion in the 10-hydroxylation reaction.

### Structural basis for 9,10-elimination activity

The experiments with DnrK RTTD F297G fortuitously led to gain-of-elimination activity (Fig. 2c) with production of **16** (15 % conversion). The active site is significantly modified by the single mutation and an explanation for 9,10-elimination may be found within the interface of R1 and R2 (Fig. 4a-4c). Key to the switch in activity resides in the fully conserved residue R285 located directly adjacent to R1 (Fig. 4c), which has gained freedom to move towards the active site in DnrK RTTD F297G. The mobility is possible due to the Y342L mutation in R2, repositioning the neighbouring D345, which removes bulky aromatic amino acids present in DnrK (Y342) and RdmB (F346) that prevent arginine-ligand interactions in the wild type enzymes.

**Figure 4.**
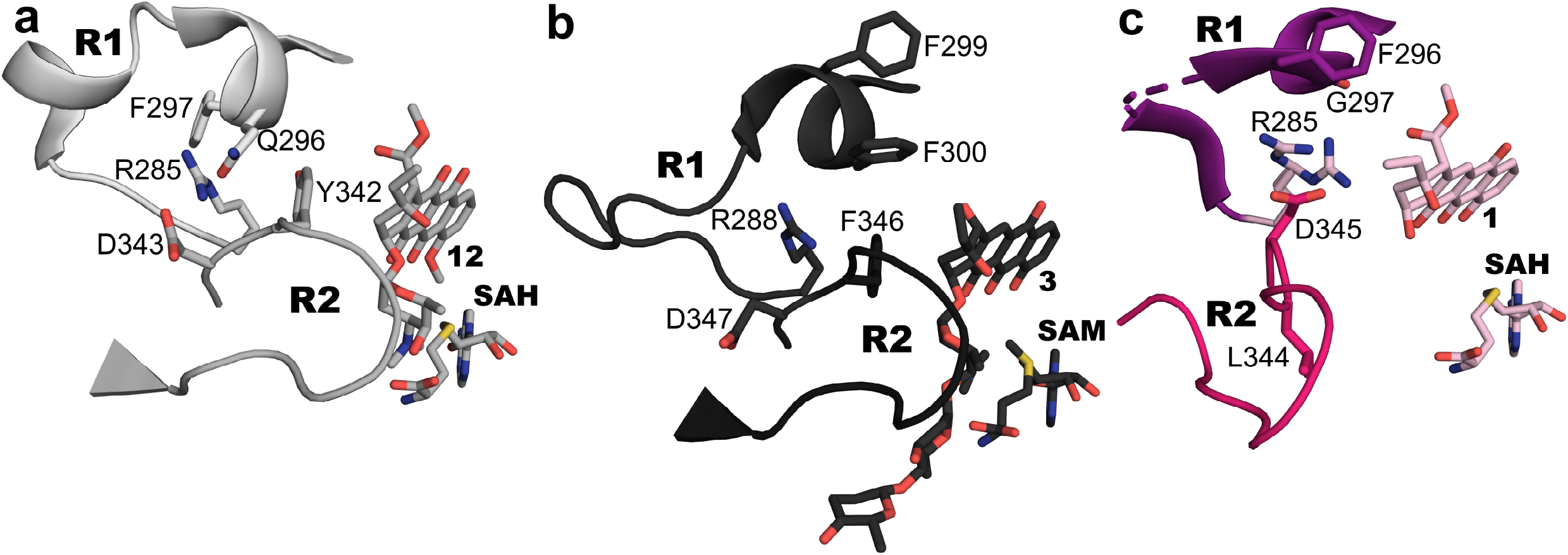
Structure elucidation of the mechanism of elimination. In the structures of **a,** DnrK (PDBID: 1TW2) and **b,** RdmB (PDBID: 1XDS) aromatic residues Y342 and F346, respectively, from R2 prevent the interaction of a conserved arginine with the substrate. **c,** DnrK RTTD F297G contains D345 in a structurally equivalent position to that of the aromatic residues Y342 in DnrK and F346 in RdmB, which provides more space for R285. Residue R285 is seen in two distinct conformations, indicating a higher degree of freedom, which will likely allow R285 to interact with the substrate.

The structural data suggests that the initial step in the 9,10-elimination reaction is 10-decarboxylation by R305 (R303 in DnrK). Formation of a carbanion intermediate is likely to be essential for the activity^31^, since DnrK RTTD F297G could not catalyse the reaction when the 10-decarboxylation product **7** was used as a substrate (Supplementary Fig. 19). Interaction of R285 with the C9 hydroxyl group of the carbanion intermediate may lead to protonation and generation of a better leaving group resulting in the elimination product.

### Factors affecting substrate specificity

TamK has been shown to catalyse 10-decarboxylation of **4** and **5**, but not **6**^29^. In order to influence reactivity towards triglycosidic substrates, we engineered R2 and R3 in the TamK scaffold. The use of amino acid sequences from RdmB had a notable synergistic effect in accepting **6**, with TamK TRRT showing the highest rate in the formation of the novel 10-decarboxylation product **14** (77 % conversion, Fig. 2e). Minor gain-of-hydroxylation activity (3 % conversion, Fig. 2e) could be seen in the construct TamK RRRT, with slightly reduced 10-decarboxylation activity (60 % conversion, Fig. 2e).

Next the chimeric pair DnrK RTTD and DnrK RTTR was constructed to examine the influence of the sugar binding regions R2 and R3 from TamK, an enzyme with higher affinity towards **4** than RdmB^29^. Both DnrK RTTD and DnrK RTTR became exceptional generalist enzymes, with the ability to use any of the substrates to catalyse reactions, yielding all potential products albeit at different yields (Fig. 2c-2e). Of particular interest is the significant gain-of-activity with the aglycone **4** in comparison to the DnrK (Fig. 2c). The presence of R1 from RdmB in both chimeras shifted the reactivity from 4-O-methylation towards 10-hydroxylation, with less than 15 % of the native DnrK activity measured, compared to almost 60 % 10-hydroxylation conversion.

Structures of DnrK RTTD and DnrK RTTR in complex with **1** revealed the molecular basis for the expanded substrate specificity. Regions R2 and R3 are observed in a closer interaction with each other, closing the active site to stabilize the smaller aglycone substrate. The modifications have altered the hydrogen bonding network between the R2 and R3 interface (Fig. 5). The α11 helix hosting R3 is extended by a single turn, which positions the helix in the space vacated by the absent carbohydrate unit. This structural change allows stabilization of the substrate via hydrogen bonding interactions between R163 and the 6-hydroxyl group of **1** (Fig. 5b and 5c).

**Figure 5.**
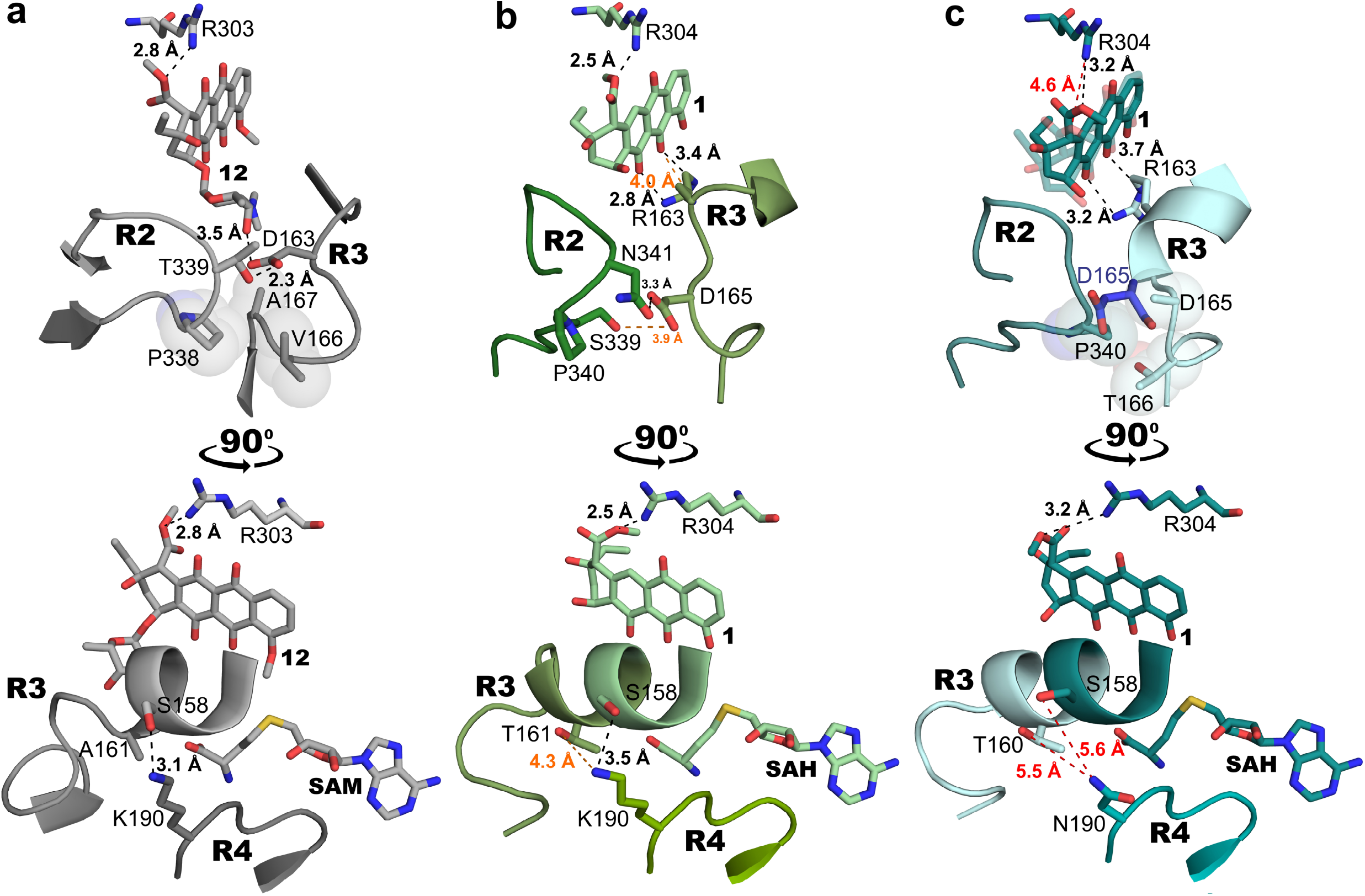
Structural basis for substrate specificity and inter-region interactions between regions R2, R3 and R4. Binding of **1** leads to rearrangements in regions R2 and R3, while the interface between regions R3 and R4 may influence substrate specificity with **6**. **a,** In DnrK (PDBID: 1TW2), D163 (R3) forms a hydrogen bond to the C3’ hydroxyl group of **12**. The interface between regions R2 and R3 shows hydrogen bonding between T339 and D163. In addition, hydrophobic interactions are present between P338 (R2) and V166 and A167 (R3). The interaction between regions R3 and R4 is due to a hydrogen bond between K190 and S158. **b,** In the absence of the sugar subunit, the extended α11 helix in DnrK RTTD positions R163 (R3) to interact with **1**. Furthermore, a more complex hydrogen-bonding network with interactions between S339 and N341 (R2) and D165 (R3) are observed. Similarly to DnrK, K190 (R4) is observed to interact weakly with residues from R3 in DnrK RTTD. **c,** The structure of DnrK RTTR shows how N190 (R4) does not interact with residues from R3, which has likely implications for the reactivity with **6**. The structure of DnrK RTTR also demonstrates mobility of non-glycosylated **1** in the active site, where the ligand is found interacting either with the conserved R304 from helix α16 (dark teal) or R163 (R3) (light teal).

In addition to R2 and R3, long-range secondary-shell effects via R4 were found to influence substrate specificity. As an example, DnrK RTTD and RTTR differ only in one amino acid, K190 and N190, respectively, that is located in R4 (Fig. 1). The activity profiles were generally similar for substrates **4** and **5**, but significant differences were observed with the triglycosylated **6** (Fig. 2). DnrK RTTD has minor total activity (8 %) with **6**, whereas DnrK RTTR is able to turn over 37 % of the substrate. The interface between R3 and R4 in DnrK in complex with **2** demonstrates that K190 from R4 is within hydrogen bonding distance (3.1 Å) to S158 located in R3 (fig. 5a). The same inter-region interaction (3.5 Å) may be observed in DnrK RTTD in complex with **1** (Fig. 5b), but the hydrogen bond is not present in DnrK RTTR in complex with **1** (Fig. 5c) due to the shorter side chain of N190 (5.6 Å). Unfolding of R3 has been observed in the complex structure of RdmB^32,40^ and appears to be essential for utilization of **6** as a substrate. The additional stabilization of helix α11 provided by the interaction of K190 and S158 in DnrK and DnrK RTTD may prevent the use of triglycosylated substrates. The proposal is further supported by DnrK RTTT, which is an efficient 10-hydroxylase for **6** (31 %) (Fig. 2). This chimera harbours P190 in the equivalent position to K190 in DnrK RTTD and hence does not exhibit strong interactions between R3 and R4 (Supplementary Fig. 20).

### The influence of region R4 and loss-of-methylation activity

The SAM binding site is highly conserved in the enzyme family with the exception of the 10-decarboxylases that contain two mutations G189A/K190P (Fig. 1). In order to investigate R4, we constructed chimera DnrK DDDT, which displayed close to no reactivity towards any substrate and led to complete loss-of-methylation activity (Fig. 2c-2e). As a control, we generated DnrK DDDR, which contains a single conservative substitution K190N and the chimera behaved similarly to the wild type enzyme. DnrK DDDR harbored no reactivity towards **4** and **6**, while retaining close to 73 % of 4-O-methylation and 11 % 10-decarboxylation activity towards **5** (Fig. 2c-2e).

These experiments lent support to the idea that the introduction of the proline may compromise correct positioning of the cofactor and quench 4-O-methylation activity. The structure of the binary complex of TamK and SAM (Supplementary Fig. 21a) confirmed the interactions between the co-substrate and R4 in 10-decarboxylases. The overall fold of TamK remains very similar to that of the other members of the family with modest root-mean-square deviations to DnrK (3.7 Å) and to RdmB (3.8 Å). However, the distinct proline-alanine pair in R4 alters the conformation of the Cα main chain in the region and leads to a movement of 2.1 Å in comparison to DnrK. This change coincides with a 1.8 Å change in SAM positioning (Supplementary Fig. 21b), which is the most likely reason behind the lack of methylation activity in TamK. Similar, but less drastic, movement can be observed also in RdmB^32^ (Supplementary Fig. 21b). The strict geometrical requirements for 4-O-methylation were also apparent in chimeras DnrK DTTD, DnrK DDCD and DnrK DCCD, where the overall activity with **5** was severely impaired, without significant gains in 4-O-methylation of **4** or **6** (Fig. 2).

### Generalist nature of the chimeric proteins

Several of the multi-chimeric proteins became truly generalist enzymes that may resemble their ancestral counterparts. DnrK RTTR can accept all substrates to carry out 10-decarboxylation, 4-O-methylation, 10-hydroxylation and 9,10-elimination (Fig. 2). The generalist nature is promoted by the mobility of **1**, which was observed in two different positions in the active site of the dimeric enzyme (Fig. 5c). In one of the monomers, the substrate is located close to the conserved R303 (DnrK numbering) critical for the 10-hydroxylation activity^32^, whereas in the other monomer the substrate has moved 1.6 Å towards R163 located in the α11 helix of R3.

The mobility of the substrates is likely to be linked to the ability of the chimeragenesis regions to adopt different conformations. An extreme example is observed in the binary complex of DnrK RTTT and **1**, where the substrate is bound in a reverse manner and partly occupies the SAM binding pocket (Supplementary Fig. 20). All chimeragenesis regions adopt significantly altered conformations to accommodate the altered binding mode of **1**, but since DnrK RTTT requires the cofactor for activity (Supplementary Fig. S22) the result may be a crystallographic artefact. Nonetheless, the significant structural changes (Supplementary Fig. 20) demonstrate the malleability of the active site, which in part can be used to explain the generalist nature of the chimeras with non-glycosylated substrates such as **4**.

Similarly, DnrK RTCR is adept at carrying out diverse reactions with improved acceptance of **6** (Fig. 2). The structure of DnrK RTCR with **2** shows that the sugar binding regions R2 and R3 no longer interact with one another (Supplementary Fig. 23), since the presence of the L-rhodosamine unit of **2** has driven R3 away from the aglycone core of the substrate. The displaced amino acids in R3 remain without a defined secondary structure, which will likely provide a higher degree of freedom for the substrate to re-arrange its position.

### Evolutionary implications

Diversification of enzyme function is fundamental to the evolution of secondary metabolism^12^. The work presented here depicts the evolutionary events that have allowed SAM-dependent methyltransferases to diversify to catalyse atypical chemical transformations. The experiments demonstrate how few point mutations in four hypervariable regions are sufficient to change protein functions (Fig. 6) and directly alter products of entire metabolic pathways.

**Figure 6.**
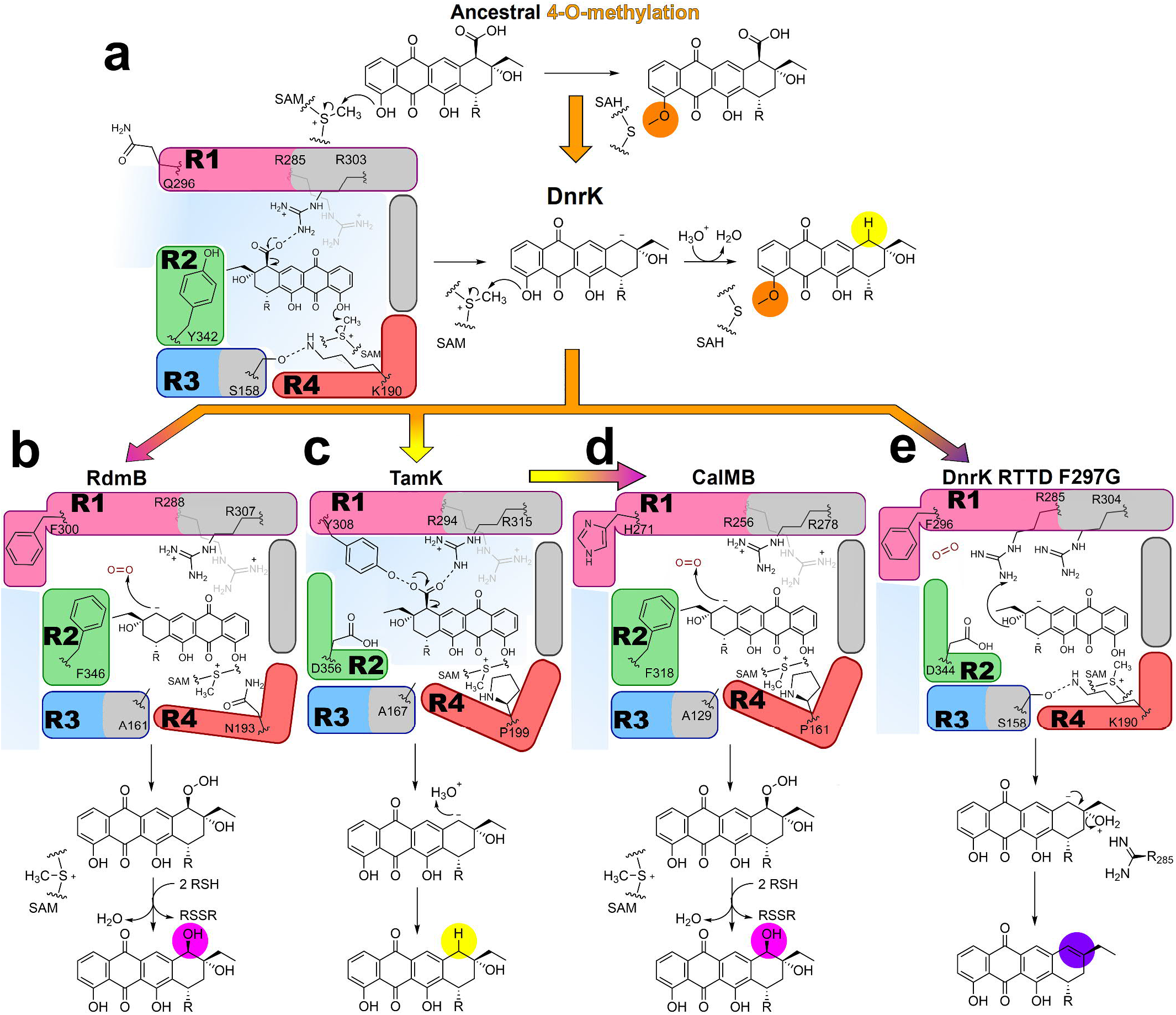
Schematic representation of different activities. Active site evolution and mechanistic consequences. **a,** The active site of 4-O-methylase DnrK shows the shared decarboxylation mechanism. In addition, its open R1 cannot prevent bulk solvent access, SAM is positioned to catalyse methyl transfer and R3 and R4 interactions prevent active site flexibility for acceptance of **6**. **b,** 10-decarboxylase TamK has Y308 interacting with the leaving decarboxyl group. R4 leads to a different binding of SAM, with no methylation. **c, d** Convergent evolution lead to two distinct 10-hydroxylases. Both CalMB (**c**) and RdmB (**d**) share an aromatic residue in R1 that closes the active site to bulk solvent and non-interacting regions R3 and R4 allowing triglycosylated anthracyclines. In the case of CalMB, R4 resembles that of TamK. **e**, The non-natural “eliminase” possesses a decarboxylase-like R2, without an aromatic residue, allowing R285 to interact with the substrate.

We propose that the initial moonlighting 10-decarboxylation activity has been the seed for functional differentiation (Fig. 6a), which has allowed the development of secondary catalytic activities^41,42^ without disturbances to 4-O-methylation. The moonlighting activity has emerged through evolution of anthracycline substrates with free 10-carboxyl groups^29,43^ in combination with conservation of R303 important for substrate binding^32,39^.

RdmB-type 10-hydroxylases (Fig. 6b) have appeared in a single evolutionary step via insertion of S298 to DnrK-type ancestral proteins^31^, leading to the repositioning of the aromatic F297 (DnrK numbering). A shared feature of DnrK and RdmB is the formation of an initial carbanion intermediate by R303. In DnrK, the intermediate reacts with H_3_O^+^ leading to 10-decarboxylation, whereas rotation of F300 and exclusion of solvent ions in RdmB allows the carbanion to react with O_2_. The hydroperoxy intermediate is stable enough to have been detected by mass spectrometry^32^ and a reaction with cellular reductants such as glutathione yields the 10-hydroxylation product.

The evolutionary events have taken a different route on the second clade of proteins. The initial step has been subfunctionalization of TamK-type proteins to catalyze solely 10-debarboxylation (Fig. 6c). Two point mutations G189A/K190P in R4 are sufficient for misalignment of SAM and loss-of-methylation activity. In addition, the 10-decarboxylation activity has been improved via changes in R1 and positioning of Y308 in a manner that enables hydrogen bonding interactions with the 10-carboxyl group (Fig. 6c).

The CalMB-type proteins present an interesting example of convergent evolution (Fig. 6d). The loss-of-methylation functionality has likely been the first step due to similarity of R4 to TamK-type proteins. Changes in R1 and particularly the Y308H (TamK numbering) mutation has led to neofunctionalization and emergence of 10-hydroxylation activity. Despite the different evolutionary paths of CalMB and RdmB, the chemistry appears to be mechanistically similar with exclusion of bulk solvent being a critical feature. The chemistry is likely to hold true also for DnrK RTTD F297G (Fig. 6e), which could be considered as a third alternative for gain-of-hydroxylation activity that Nature has not yet discovered.

Critical for the laboratory evolution of the novel 9,10-elimination activity appears to be the removal of bulky aromatic amino acids in R2 (Fig. 6e). Mutations such as Y342L in DnrK RTTD F297G allow repositioning of a fully conserved R285 (DnrK numbering) towards the active site and interaction with the substrate (Fig. 4). The 9,10-elimination is likely to proceed via the same carbanion intermediate that is also important for 10-decarboxylation and 10-hydroxylation activities (Fig. 6b-6d). The artificial evolution of 9,10-elimination functionality is encouraging as it demonstrates that the catalytic repertoire of natural product biosynthesis enzymes can be further engineered in the laboratory.

## Methods

### Reagents

All reagents were bought from SigmaAldrich unless stated otherwise. TALON SuperFlow resin and PD-10 desalting columns were bought from GE Healthcare. SDS-PAGE Bolt™ 4-12% Bis-Tris Plus Gel and PageRuler™ Unstained Protein Ladder molecular weight standard were bought from Thermo Fisher Scientific. Phusion polymerase, restriction enzymes, T4 DNA ligases, and GeneJET Plasmid Miniprep Kit were bought from Thermo Fisher Scientific.

### Production and purification of compounds

Compounds **1, 2**, and **3** were produced and purified as described previously^29,31^. Briefly, aclacinomycin T (**2**) was obtained from a fermentation of strain *Streptomyces galilaeus* ATCC 31615 mutant H038^44^. Aclacinomycin A (**3**) was obtained in a two-step fermentation process by first cultivating *Streptomyces galilaeus* ATCC 31615 mutant HO42 for production of aclacinomycin B, followed by biotransformation to **3** using Streptomyces galilaeus ATCC 31615 mutant HO26^44^. Aklavinone (**1**) was obtained through acid hydrolysis of **3** by using equal volumes of toluene and 2.5 M HCl at 60 °C for 1 h 20 min. The compounds were purified by preparative HPLC using a LC-20AP/CBM-20A system with a diode array detector (Shimadzu) and EVO C18, 5 μm, 100 Å, 250 x 21.2 mm Kinetex column (Phenomenex). Purity of the compounds were estimated by HPLC and was at least 96 % for **1**, 88 % for **2** and 76 % for **3**.

### Protein expression and purification

Protein expression and purification were done as described previously^29,31^. Briefly, protein expression was carried out in *Escherichia coli* TOP10 cells. Protein purification was carried out through a polyHis-tag affinity chromatography with TALON SuperFlow resin and PD-10 desalting columns. Enzymes were concentrated using Amicon Ultra 0.5 mL centrifugation filters (10 000 nominal molecular weight limit). Purity of all the enzymes was estimated to be 90 % or more by SDS-PAGE.

### General DNA Techniques

Modified pBAD/His B vector (Invitrogen) containing native *dnrK* and *tamK*^31,39^ were used as templates for chimeragenesis. The DrnK chimeras TDDD, CDDD, DTTD, RTTD, RTTR, and TamK chimeras DRRD, RRTT, DRTD, DDRD, TRRT, DDTD, RRRT, DDDR and DDDD were ordered as synthetic DNA from GeneArt, Thermo Fisher or Sigma Aldrich (Strings DNA Fragments). The DnrK chimeras DDDT, DDCD and DCCD were generated using the four-primer overhang extension method as described previously^31^. In addition, both synthetic DNA and four-primer overhang extension method were utilized to construct the following DnrK chimeras: RTTT, RTCR, RCCR and RTTD F297G. The obtained chimeras were cloned into pBAD/His B vectors. All DNA sequences were confirmed by sequencing before protein expression.

### Enzyme Activity Measurements

Enzymatic activity measurements were carried out as previously described^29^. Briefly, a two-step reaction was initiated by incubation of **1** (120 μM) with EamC (9 μM) or alternatively **2** (120 μM) or **3** (120 μM) with DnrP (130 μM), to carry out an initial 15-methyl removal. The reaction products of this initial step were isolated as described previously^31^. Activity measurements of the chimeric proteins were performed with the 15-demethylated compounds **4, 5**, and **6** under the following conditions: 100 mM Tris·HCl (pH 7.5), 10 mM DTT and 400 μM SAM. The concentration of the native and chimeric enzymes was set to 6 μM.

All reactions were monitored by HPLC (SCL-10Avp/SpdM10Avp system with a diode array detector (Shimadzu)) using a C18 column (2.6 μm, 100 Å, 4.6 × 100 mm Kinetex column (Phenomenex)). The gradient elution used consisted of an initial aqueous buffer (5 % (vol/vol) acetonitrile with 45 % ammonium acetate - acetic acid (pH 3.6)) that was gradually changed to an elution buffer (100 % acetonitrile). All compounds reported were confirmed by low-resolution MS (Agilent 6120 Quadrupole LCMS system; linked to an Agilent Technologies 1260 infinity HPLC system) with identical columns, gradient, and buffer systems as described previously^31^.

### HR-MS and NMR analysis of novel compounds

Analysis by HR-MS of **13, 15**, and **16** was carried out by dissolving the compound in methanol, followed by direct injection to MicrOTOF-Q high resolution MS (Bruker Daltonics). A sample of the same compound **16** was desiccated overnight, dissolved in deuterated chloroform and subjected to NMR analyses. The spectra were recorded with a 500 MHz instrument (Bruker) with a liquid nitrogen cooled Prodigy BBO (CryoProbe), a 600 MHz instrument (Bruker) with a liquid nitrogen cooled Prodigy TCI (inverted CryoProbe) at 298 K. The signals were internally referenced to the solvent signals or tetramethylsilane. The experiments included 1D spectral analysis (1H, 13C) and 2D measurements (COSY, HMBC, and HSQCDE). Topspin (Bruker Biospin) was used for spectral analysis.

### Crystallization, Data Collection and Structure Determination

Purified proteins were mixed with both the relevant anthracycline (100 mM stock in DMSO) and SAM (50 mM stock in buffer) to a 1:3:3 molar stoichiometry of protein:anthracycline:SAM, to a final protein concentration of 25 mg/mL. Crystals were set-up via the hanging drop method in a 1:1 ratio with the crystallization condition.

Crystals of DnrK RTTD, DnrK RTTR and DnrK RTTR F297G were obtained in 0.1 M Sodium acetate, pH 4.6; 8 % PEG 8000, by co-crystallization with **1**. Attempts to co-crystallize with **2** yielded data with an aglycone present in the active site, likely caused by the acidic cleavage of the O-glycosidic bond.

DnrK RTCR crystallized in 0.2 M Magnesium chloride hexahydrate; 0.1 M HEPES 7.0; 20 *% w/v* PEG 6000, in the presence of **2**. DnrK RTTT crystallized under the same condition, with varying concentrations of SAM, however no density for the co-factor was observed in the active site. The samples were incubated for 6 days at 21 °C for crystal growth. DnrK TDDD crystallized in 0.2 M Magnesium chloride hexahydrate; 0.1 M Tris-HCl (pH=8.0); 20 % w/v PEG 6000 in the presence of **2.** The samples were incubated for 14 days at 4 °C for crystal growth. DnrK CDDD crystallized in 0.2 M Potassium thiocyanate, 0.1 M Bis-Tris propane (pH= 7.5), 20 % w/v PEG 3350 in the presence of **2**. The samples were incubated for 21 days at 4 °C for crystal growth. After crystal formation the samples were collected and flash frozen in liquid nitrogen.

Diffraction data for DnrK RTTD, DnrK RTTR, DnrK RTTR F297G, DnrK RTCR and DnrK RTTT were collected at the Max IV facilities (Lund, Sweden) at the BioMax beamline. Diffraction data for DnrK TDDD and DnrK CDDD was collected at the European Synchrotron Radiation Facilities (ESRF) (Grenoble, France), at the ID30A-1 beamline. Details of the data collection and refinement statistics are listed in Supplementary Table 3. Data were indexed, integrated and scaled in XDS^45^ and AIMLESS^46^. Due to anisotropic diffraction in the DnrK RTTR dataset, ellipsoidal truncation followed by anisotropic scaling was carried out in the UCLA-DOE LAB Diffraction Anisotropy Server^47^. Initial phases were obtained by molecular replacement with the DnrK WT structure (1tw2) as a starting model, using the Phenix Molecular Replacement program (Phaser)^48^. Figures depicting protein structures were prepared using PyMOL (The PyMOL Molecular Graphics System, version 1.3; Schrödinger, LLC).

