## Supporting information for "Evolution-Inspired Engineering of Anthracycline Methyltransferases"

**Supplementary information**

**Supplementary Table 1. Chimeric sequences being studied.** Amino acid sequences of the enzyme chimeras under study are presented, with regions under study in bold. The single point mutation introduced in the chimera DnrK RTTD F297G is indicated in red.

| **Protein name** | **Amino acid sequence** |
| --- | --- |
| DnrK WT | MAHHHHHHHRSTAEPTVAARPQQIDALRTLIRLGSLHTPMVV RTAATLRLVDHILAGARTVKALAARTDTRPEALLRLIRHLVA IGLLEEDAPGEFVPTEVGELLADDHPAAQRAWHDLTQAVARA DISFTRLPDAIRTGRPTYESIYGKPFYEDLAGRPDLRASFDS L**LACDQDVAFD**APAAAYDWTNVRHVL**DVGGGKG**GFAAAIARR APHVSATVLEMAGTVDTARSYLKDEGLSDRVDVVEGDFFEPL PRKADAIILSFVLLNWPDHDAVRILTRCAEALEPGGRILIHE R**DDLHENSFNEQFTE**LDLRMLVFLGGALRTREKWDGLAASAG LVVEEVR**QLPSPTIPYDLS**LLVLAPAATGA |
| DnrK RDDD | MAHHHHHHHRSTAEPTVAARPQQIDALRTLIRLGSLHTPMVV RTAATLRLVDHILAGARTVKALAARTDTRPEALLRLIRHLVA IGLLEEDAPGEFVPTEVGELLADDHPAAQRAWHDLTQAVARA DISFTRLPDAIRTGRPTYESIYGKPFYEDLAGRPDLRASFDS LLACDQDVAFDAPAAAYDWTNVRHVLDVGGGKGGFAAAIARR APHVSATVLEMAGTVDTARSYLKDEGLSDRVDVVEGDFFEPL PRKADAIILSFVLLNWPDHDAVRILTRCAEALEPGGRILIHE R**ADVEGDGADRFFSTL**LDLRMLVFLGGALRTREKWDGLAASA GLVVEEVRQLPSPTIPYDLSLLVLAPAATGA |
| DnrK CDDD | MAHHHHHHHRSTAEPTVAARPQQIDALRTLIRLGSLHTPMVV RTAATLRLVDHILAGARTVKALAARTDTRPEALLRLIRHLVA IGLLEEDAPGEFVPTEVGELLADDHPAAQRAWHDLTQAVARA DISFTRLPDAIRTGRPTYESIYGKPFYEDLAGRPDLRASFDS LLACDQDVAFDAPAAAYDWTNVRHVLDVGGGKGGFAAAIARR APHVSATVLEMAGTVDTARSYLKDEGLSDRVDVVEGDFFEPL PRKADAIILSFVLLNWPDHDAVRILTRCAEALEPGGRILIHE R**AEPSPDETTSTADLHFSL**LDLRMLVFLGGALRTREKWDGLA ASAGLVVEEVRQLPSPTIPYDLSLLVLAPAATGA |
| DnrK TDDD | MAHHHHHHHRSTAEPTVAARPQQIDALRTLIRLGSLHTPMVV RTAATLRLVDHILAGARTVKALAARTDTRPEALLRLIRHLVA IGLLEEDAPGEFVPTEVGELLADDHPAAQRAWHDLTQAVARA DISFTRLPDAIRTGRPTYESIYGKPFYEDLAGRPDLRASFDS LLACDQDVAFDAPAAAYDWTNVRHVLDVGGGKGGFAAAIARR APHVSATVLEMAGTVDTARSYLKDEGLSDRVDVVEGDFFEPL PRKADAIILSFVLLNWPDHDAVRILTRCAEALEPGGRILIHE R**AEAPSGGTRTSDLYFSV**LDLRMLVFLGGALRTREKWDGLAA SAGLVVEEVRQLPSPTIPYDLSLLVLAPAATGA |
| DnrK DDDT | MAHHHHHHHRSTAEPTVAARPQQIDALRTLIRLGSLHTPMVV RTAATLRLVDHILAGARTVKALAARTDTRPEALLRLIRHLVA IGLLEEDAPGEFVPTEVGELLADDHPAAQRAWHDLTQAVARA DISFTRLPDAIRTGRPTYESIYGKPFYEDLAGRPDLRASFDS LLACDQDVAFDAPAAAYDWTNVRHVL**DVGGAPG**GFAAAIARR APHVSATVLEMAGTVDTARSYLKDEGLSDRVDVVEGDFFEPL PRKADAIILSFVLLNWPDHDAVRILTRCAEALEPGGRILIHE RDDLHENSFNEQFTELDLRMLVFLGGALRTREKWDGLAASAG LVVEEVRQLPSPTIPYDLSLLVLAPAATGA |
| DnrK DDDR | MAHHHHHHHRSTAEPTVAARPQQIDALRTLIRLGSLHTPMVV RTAATLRLVDHILAGARTVKALAARTDTRPEALLRLIRHLVA IGLLEEDAPGEFVPTEVGELLADDHPAAQRAWHDLTQAVARA DISFTRLPDAIRTGRPTYESIYGKPFYEDLAGRPDLRASFDS LLACDQDVAFDAPAAAYDWTNVRHVL**DVGGGNG**GFAAAIARR APHVSATVLEMAGTVDTARSYLKDEGLSDRVDVVEGDFFEPL PRKADAIILSFVLLNWPDHDAVRILTRCAEALEPGGRILIHE RDDLHENSFNEQFTELDLRMLVFLGGALRTREKWDGLAASAG LVVEEVRQLPSPTIPYDLSLLVLAPAATGA |
| DnrK DDCD | MAHHHHHHHRSTAEPTVAARPQQIDALRTLIRLGSLHTPMVV RTAATLRLVDHILAGARTVKALAARTDTRPEALLRLIRHLVA IGLLEEDAPGEFVPTEVGELLADDHPAAQRAWHDLTQAVARA DISFTRLPDAIRTGRPTYESIYGKPFYEDLAGRPDLRASFDS L**MATEEEAVDE**APAAAYDWTNVRHVLDVGGGKGGFAAAIARR APHVSATVLEMAGTVDTARSYLKDEGLSDRVDVVEGDFFEPL PRKADAIILSFVLLNWPDHDAVRILTRCAEALEPGGRILIHE RDDLHENSFNEQFTELDLRMLVFLGGALRTREKWDGLAASAG LVVEEVRQLPSPTIPYDLSLLVLAPAATGA |
| DnrK DCCD | MAHHHHHHHRSTAEPTVAARPQQIDALRTLIRLGSLHTPMVV RTAATLRLVDHILAGARTVKALAARTDTRPEALLRLIRHLVA IGLLEEDAPGEFVPTEVGELLADDHPAAQRAWHDLTQAVARA DISFTRLPDAIRTGRPTYESIYGKPFYEDLAGRPDLRASFDS L**MATEEEAVDE**APAAAYDWTNVRHVLDVGGGKGGFAAAIARR APHVSATVLEMAGTVDTARSYLKDEGLSDRVDVVEGDFFEPL PRKADAIILSFVLLNWPDHDAVRILTRCAEALEPGGRILIHE RDDLHENSFNEQFTELDLRMLVFLGGALRTREKWDGLAASAG LVVEEVR**PITSPVVPFDFC**LLVLAPAATGA |
| DnrK DTTD | MAHHHHHHHRSTAEPTVAARPQQIDALRTLIRLGSLHTPMVV RTAATLRLVDHILAGARTVKALAARTDTRPEALLRLIRHLVA IGLLEEDAPGEFVPTEVGELLADDHPAAQRAWHDLTQAVARA DISFTRLPDAIRTGRPTYESIYGKPFYEDLAGRPDLRASFDS L**MTTREDTAFA**APAAAYDWTNVRHVLDVGGGKGGFAAAIARR APHVSATVLEMAGTVDTARSYLKDEGLSDRVDVVEGDFFEPL PRKADAIILSFVLLNWPDHDAVRILTRCAEALEPGGRILIHE RDDLHENSFNEQFTELDLRMLVFLGGALRTREKWDGLAASAG LVVEEVR**GPLVSPNVPLDSC**LLVLAPAATGA |
| DnrK RTTD | MAHHHHHHHRSTAEPTVAARPQQIDALRTLIRLGSLHTPMVV RTAATLRLVDHILAGARTVKALAARTDTRPEALLRLIRHLVA IGLLEEDAPGEFVPTEVGELLADDHPAAQRAWHDLTQAVARA DISFTRLPDAIRTGRPTYESIYGKPFYEDLAGRPDLRASFDS L**MTTREDTAFA**APAAAYDWTNVRHVLDVGGGKGGFAAAIARR APHVSATVLEMAGTVDTARSYLKDEGLSDRVDVVEGDFFEPL PRKADAIILSFVLLNWPDHDAVRILTRCAEALEPGGRILIHE R**ADVEGDGADRFFSTL**LDLRMLVFLGGALRTREKWDGLAASA GLVVEEVR**GPLVSPNVPLDSC**LLVLAPAATGA |
| DnrK RTTD F297G | MAHHHHHHHRSTAEPTVAARPQQIDALRTLIRLGSLHTPMVV RTAATLRLVDHILAGARTVKALAARTDTRPEALLRLIRHLVA IGLLEEDAPGEFVPTEVGELLADDHPAAQRAWHDLTQAVARA DISFTRLPDAIRTGRPTYESIYGKPFYEDLAGRPDLRASFDS L**MTTREDTAFA**APAAAYDWTNVRHVLDVGGGKGGFAAAIARR APHVSATVLEMAGTVDTARSYLKDEGLSDRVDVVEGDFFEPL PRKADAIILSFVLLNWPDHDAVRILTRCAEALEPGGRILIHE R**ADVEGDGADRFGSTL**LDLRMLVFLGGALRTREKWDGLAASA GLVVEEVR**GPLVSPNVPLDSC**LLVLAPAATGA |
| DnrK RTTR | MAHHHHHHHRSTAEPTVAARPQQIDALRTLIRLGSLHTPMVV RTAATLRLVDHILAGARTVKALAARTDTRPEALLRLIRHLVA IGLLEEDAPGEFVPTEVGELLADDHPAAQRAWHDLTQAVARA DISFTRLPDAIRTGRPTYESIYGKPFYEDLAGRPDLRASFDS L**MTTREDTAFA**APAAAYDWTNVRHVL**DVGGGNG**GFAAAIARR APHVSATVLEMAGTVDTARSYLKDEGLSDRVDVVEGDFFEPL PRKADAIILSFVLLNWPDHDAVRILTRCAEALEPGGRILIHE R**ADVEGDGADRFFSTL**LDLRMLVFLGGALRTREKWDGLAASA GLVVEEVR**GPLVSPNVPLDSC**LLVLAPAATGA |
| DnrK RTCR | MAHHHHHHHRSTAEPTVAARPQQIDALRTLIRLGSLHTPMVV RTAATLRLVDHILAGARTVKALAARTDTRPEALLRLIRHLVA IGLLEEDAPGEFVPTEVGELLADDHPAAQRAWHDLTQAVARA DISFTRLPDAIRTGRPTYESIYGKPFYEDLAGRPDLRASFDS L**MATEEEAVDE**APAAAYDWTNVRHVL**DVGGGNG**GFAAAIARR APHVSATVLEMAGTVDTARSYLKDEGLSDRVDVVEGDFFEPL PRKADAIILSFVLLNWPDHDAVRILTRCAEALEPGGRILIHE R**ADVEGDGADRFFSTL**LDLRMLVFLGGALRTREKWDGLAASA GLVVEEVR**GPLVSPNVPLDSC**LLVLAPAATGA |
| DnrK RCCR | MAHHHHHHHRSTAEPTVAARPQQIDALRTLIRLGSLHTPMVV RTAATLRLVDHILAGARTVKALAARTDTRPEALLRLIRHLVA IGLLEEDAPGEFVPTEVGELLADDHPAAQRAWHDLTQAVARA DISFTRLPDAIRTGRPTYESIYGKPFYEDLAGRPDLRASFDS L**MATEEEAVDE**APAAAYDWTNVRHVL**DVGGGNG**GFAAAIARR APHVSATVLEMAGTVDTARSYLKDEGLSDRVDVVEGDFFEPL PRKADAIILSFVLLNWPDHDAVRILTRCAEALEPGGRILIHE R**ADVEGDGADRFFSTL**LDLRMLVFLGGALRTREKWDGLAASA GLVVEEVR**PITSPVVPFDFC**LLVLAPAATGA |
| DnrK RTTT | MAHHHHHHHRSTAEPTVAARPQQIDALRTLIRLGSLHTPMVV RTAATLRLVDHILAGARTVKALAARTDTRPEALLRLIRHLVA IGLLEEDAPGEFVPTEVGELLADDHPAAQRAWHDLTQAVARA DISFTRLPDAIRTGRPTYESIYGKPFYEDLAGRPDLRASFDS L**MTTREDTAFA**APAAAYDWTNVRHVL**DVGGAPG**GFAAAIARR APHVSATVLEMAGTVDTARSYLKDEGLSDRVDVVEGDFFEPL PRKADAIILSFVLLNWPDHDAVRILTRCAEALEPGGRILIHE R**ADVEGDGADRFFSTL**LDLRMLVFLGGALRTREKWDGLAASA GLVVEEVR**GPLVSPNVPLDSC**LLVLAPAATGA |
| TamK WT | MAHHHHHHHRSSGTDAGTAGTAGTAGAGAGGDRQHVDALVRM SNLVTPMALRVAATLRLVDHLRAGATSADALADATGADADAL ARLMRHLAAAGVLEEPEPGHYAPTGLGDLLADDHPSRQRSWL DLDQAVGRADLTFLGLREAVRTGRPQYEARYGKPFWTDLSED DGLGASFDAL**MTTREDTAFA**APVAAYDWTRARHVL**DVGGAPG** GLLTAILRAAPEAHGTLLDLPGAAARTRERIAANGMDERIDV VGGDFFDELPVTADVVVLSFTLLNWSDPDALRILGRCRDALR PGGRIVLLER**AEAPSGGTRTSDLYFSV**LDMRMLVFLGGRVRT DREWADLAAAAGLDIVGKT**GPLVSPNVPLDSC**LWELAPR |
| TamK RRTT | MAHHHHHHHRSSGTDAGTAGTAGTAGAGAGGDRQHVDALVRM SNLVTPMALRVAATLRLVDHLRAGATSADALADATGADADAL ARLMRHLAAAGVLEEPEPGHYAPTGLGDLLADDHPSRQRSWL DLDQAVGRADLTFLGLREAVRTGRPQYEARYGKPFWTDLSED DGLGASFDALMTTREDTAFAAPVAAYDWTRARHVLDVGGAPG GLLTAILRAAPEAHGTLLDLPGAAARTRERIAANGMDERIDV VGGDFFDELPVTADVVVLSFTLLNWSDPDALRILGRCRDALR PGGRIVLLER**ADVEGDGADRFFSTL**LDMRMLVFLGGRVRTDR EWADLAAAAGLDIVGKTG**SGSTTLPFDFS**LWELAPR |
| TamK TRRT | MAHHHHHHHRSSGTDAGTAGTAGTAGAGAGGDRQHVDALVRM SNLVTPMALRVAATLRLVDHLRAGATSADALADATGADADAL ARLMRHLAAAGVLEEPEPGHYAPTGLGDLLADDHPSRQRSWL DLDQAVGRADLTFLGLREAVRTGRPQYEARYGKPFWTDLSED DGLGASFDAL**MSCDEDLAYE**APVAAYDWTRARHVLDVGGAPG GLLTAILRAAPEAHGTLLDLPGAAARTRERIAANGMDERIDV VGGDFFDELPVTADVVVLSFTLLNWSDPDALRILGRCRDALR PGGRIVLLERAEAPSGGTRTSDLYFSVLDMRMLVFLGGRVRT DREWADLAAAAGLDIVGKTG**SGSTTLPFDFS**LWELAPR |
| TamK RRRT | MAHHHHHHHRSSGTDAGTAGTAGTAGAGAGGDRQHVDALVRM SNLVTPMALRVAATLRLVDHLRAGATSADALADATGADADAL ARLMRHLAAAGVLEEPEPGHYAPTGLGDLLADDHPSRQRSWL DLDQAVGRADLTFLGLREAVRTGRPQYEARYGKPFWTDLSED DGLGASFDAL**MSCDEDLAYE**APVAAYDWTRARHVLDVGGAPG GLLTAILRAAPEAHGTLLDLPGAAARTRERIAANGMDERIDV VGGDFFDELPVTADVVVLSFTLLNWSDPDALRILGRCRDALR PGGRIVLLER**ADVEGDGADRFFSTL**LDMRMLVFLGGRVRTDR EWADLAAAAGLDIVGKTG**SGSTTLPFDFS**LWELAPR |

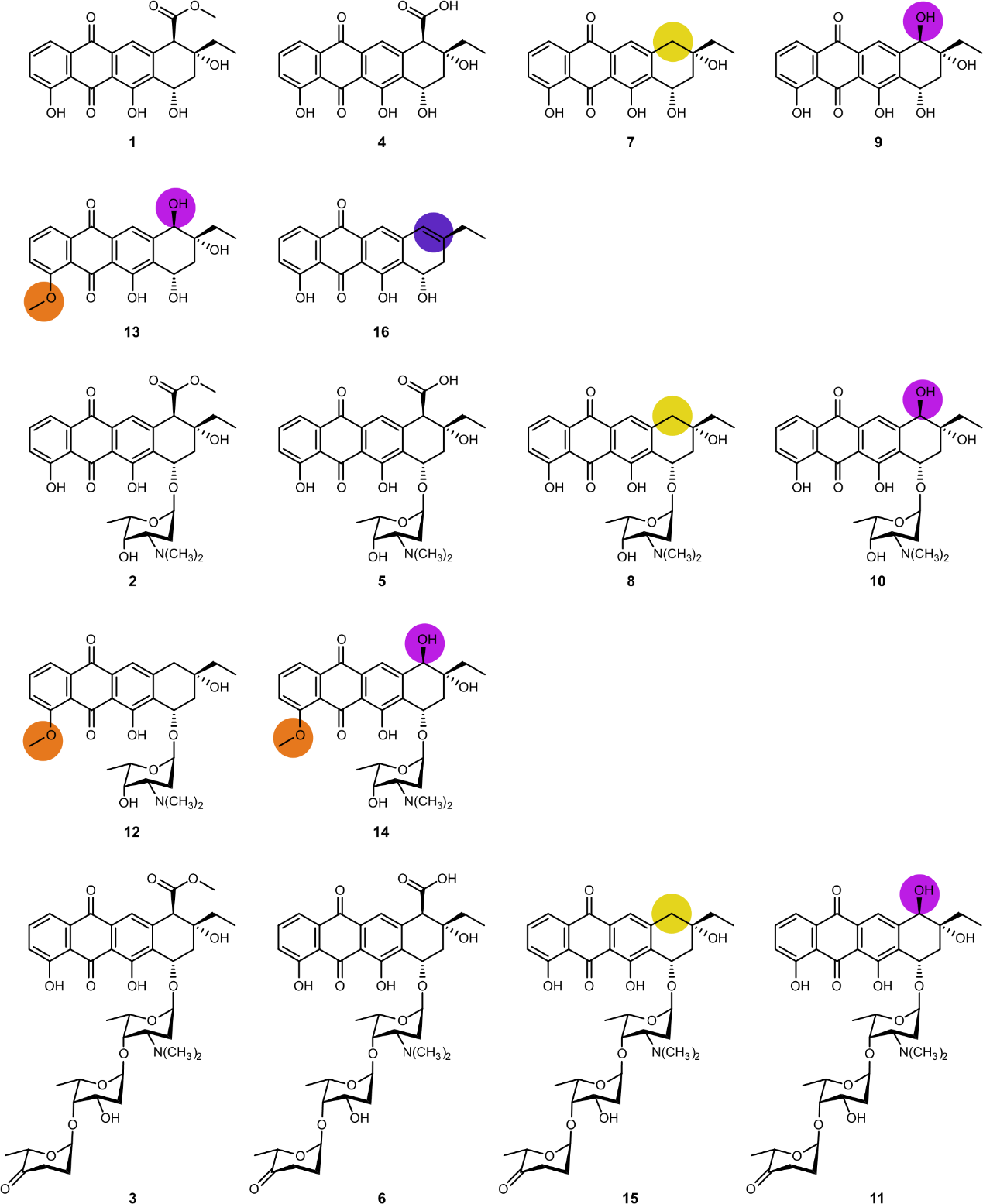

**Supplementary Figure 1.** **All the chemical structures presented in this study.**

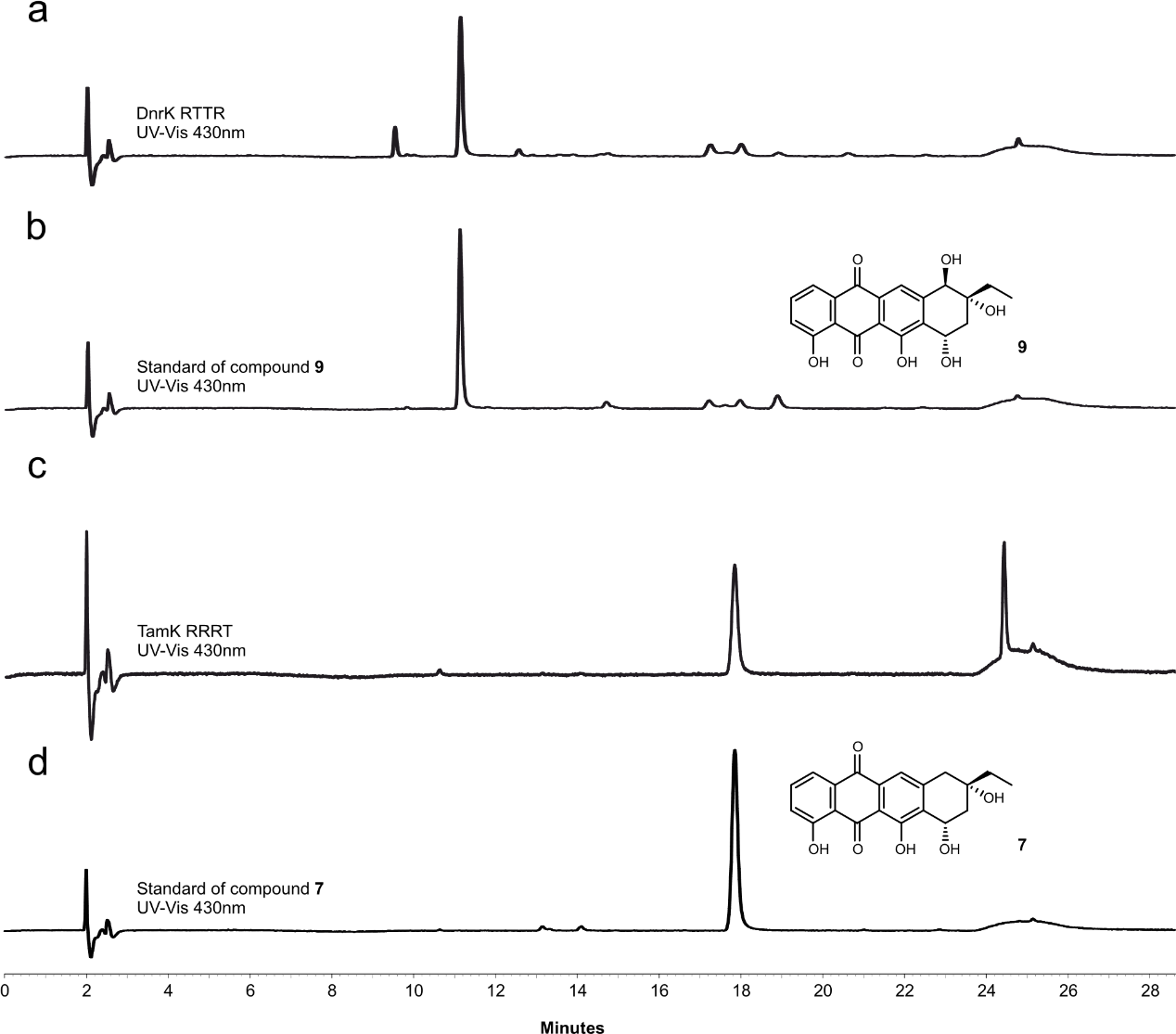

**Supplementary Figure 2.** **HPLC comparisons of enzymatic reactions products from 4 as a substrate with standard compounds.** UV-Vis chromatogram traces were recorded at 430 nm. **a,** Enzymatic reaction products by DnrK RTTR. **b,** Standard of compound **9** obtained by enzymatic reaction of RdmB with **4** as a substrate. **c,** Enzymatic reaction products by TamK RRRT. **d,** Standard of compound **7** obtained by enzymatic reaction of TamK with **4** as a substrate.

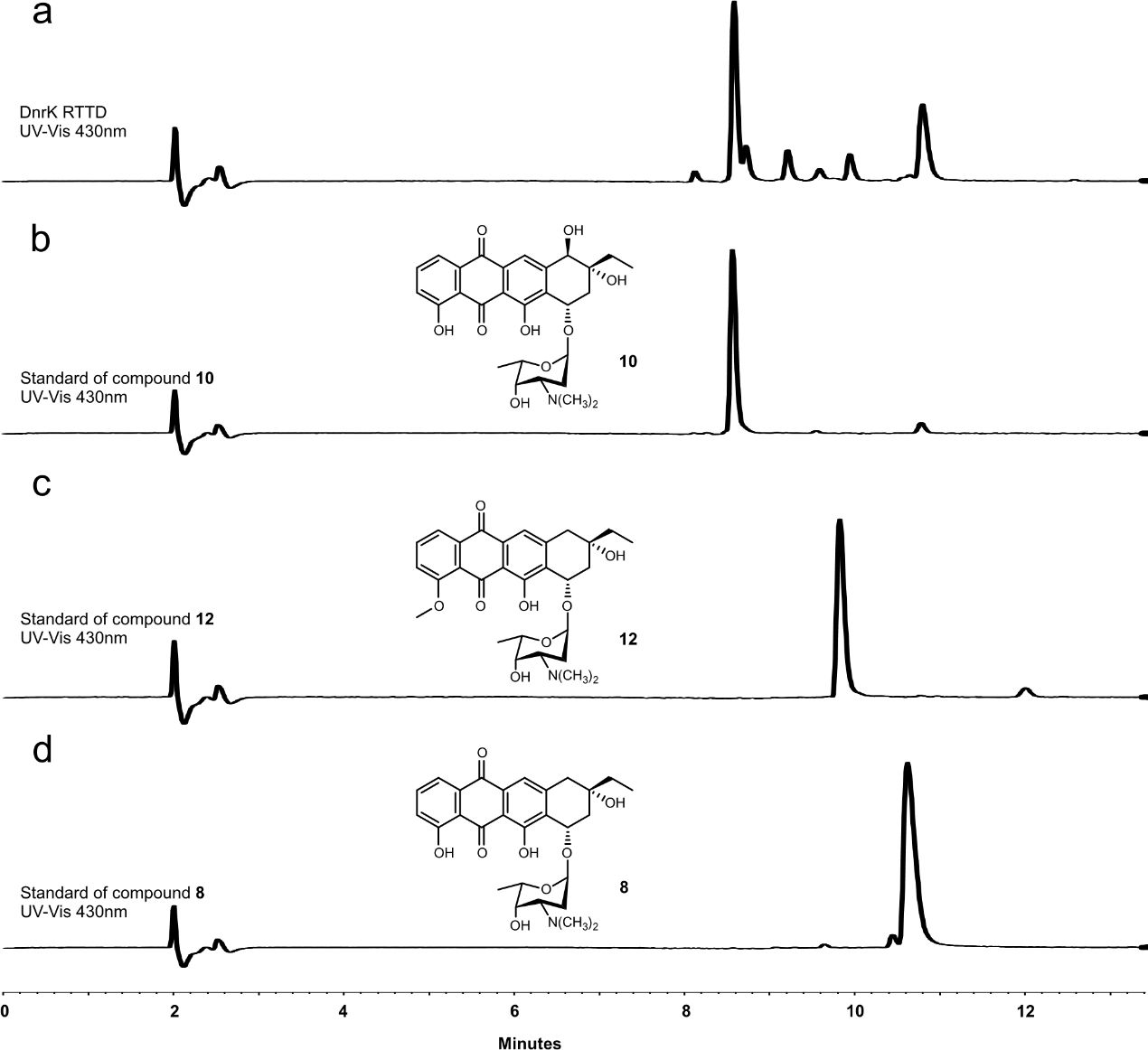

**Supplementary Figure 3.** **HPLC comparisons of enzymatic reactions products from 5 as a substrate with standard compounds.** UV-Vis chromatogram traces were recorded at 430 nm. **a,** Enzymatic reaction products by DnrK RTTD. **b,** Standard of compound **10** obtained by enzymatic reaction of RdmB with **5** as a substrate. **c,** Standard of compound **12** obtained by enzymatic reaction of DnrK with **5** as a substrate. **d,** Standard of compound **8** obtained by enzymatic reaction of TamK with **5** as a substrate.

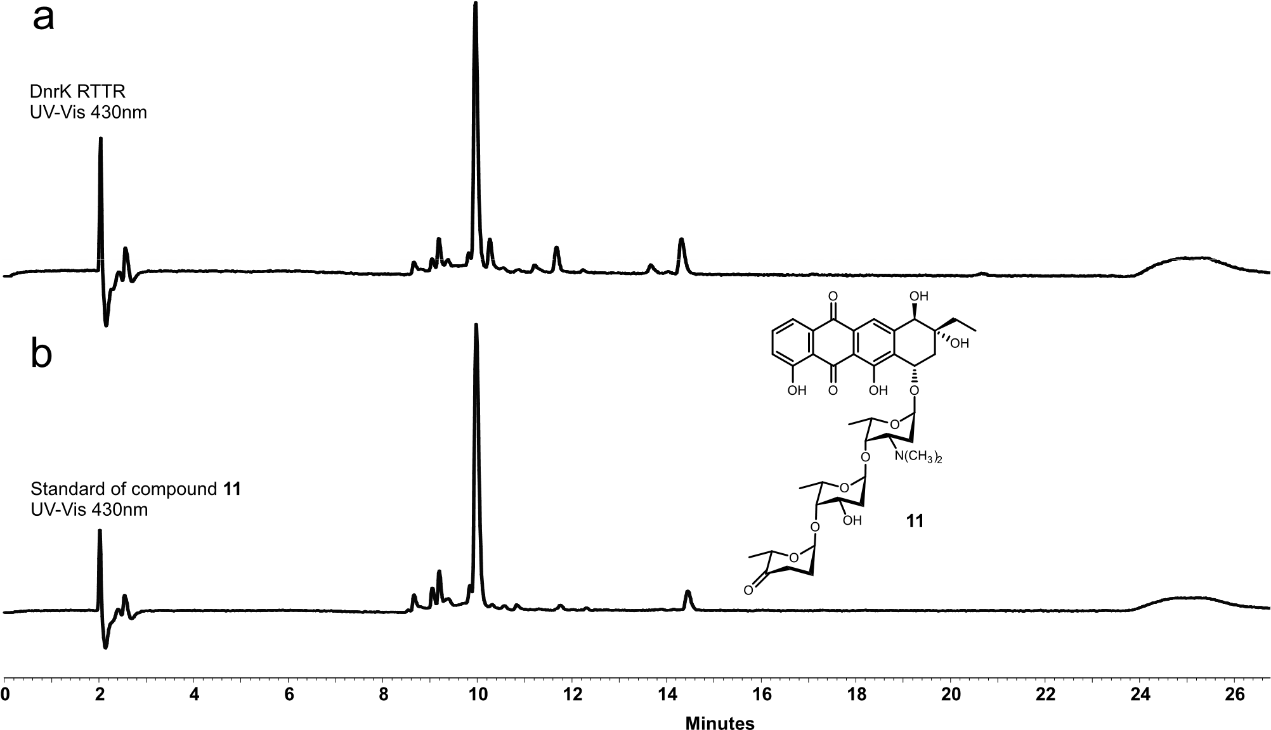

**Supplementary Figure 4.** **HPLC comparisons of enzymatic reactions products from 6 as a substrate with standard compounds**. UV-Vis chromatogram traces were recorded at 430 nm. **a,** Enzymatic reaction products by DnrK RTTR. **b,** Standard of compound **11** obtained by enzymatic reaction of RdmB with **6** as a substrate.

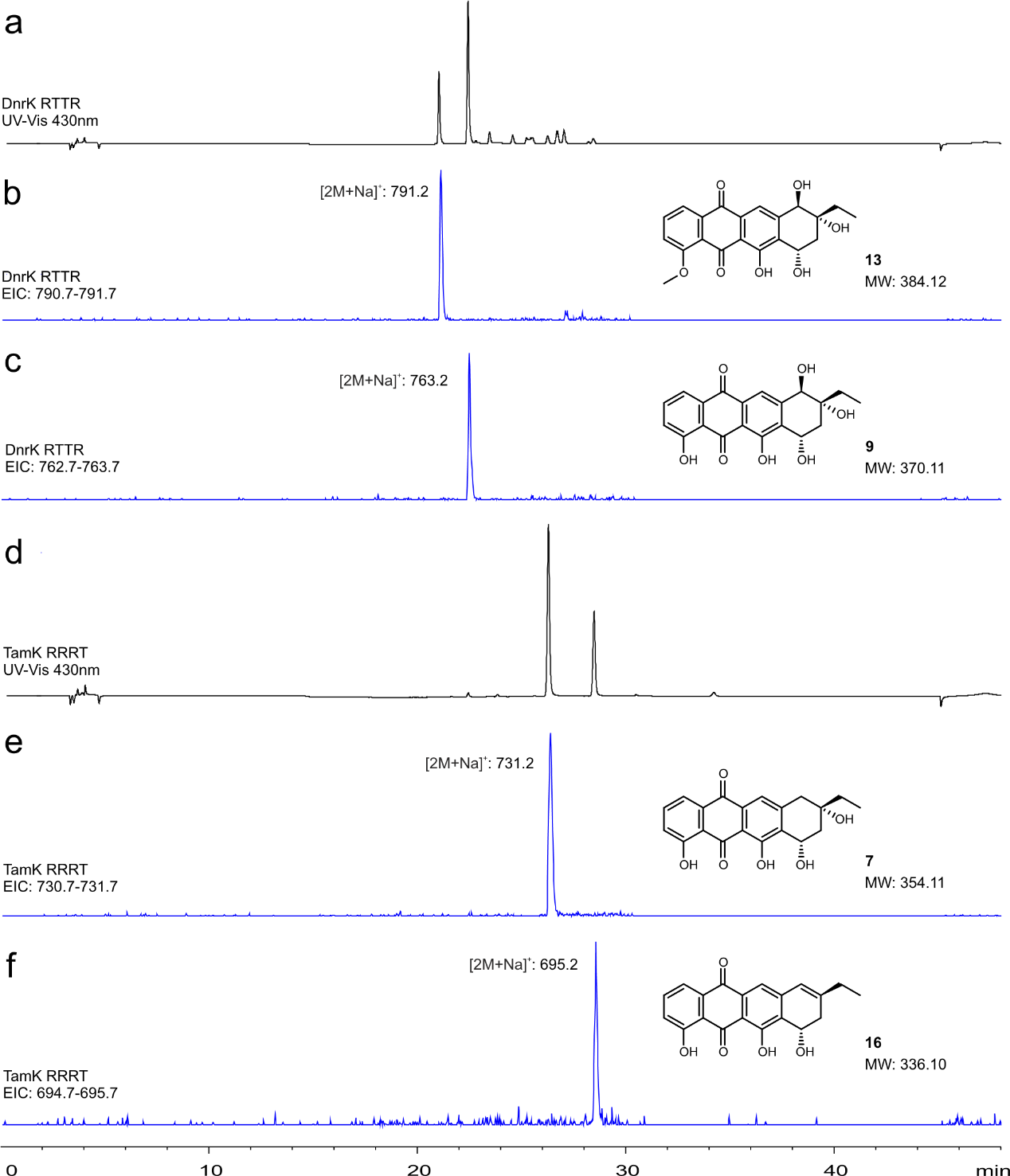

**Supplementary Figure 5. LC-MS analysis of enzymatic reactions with 4 as a substrate. a,** UV-Vis chromatogram trace recorded at 430 nm for enzymatic reaction products of DnrK RTTR. **b** **and** **c,** Extracted ion chromatogram traces in positive mode for DnrK RTTR products. **d,** UV-Vis chromatogram trace recorded at 430 nm for enzymatic reaction products of TamK RRRT. **e** **and** **f,** Extracted ion chromatogram traces in positive mode for TamK RRRT products. Products are observed as sodium adducts [2M+Na]^+^ under the conditions used, giving consistently values of 2M + 22.99.

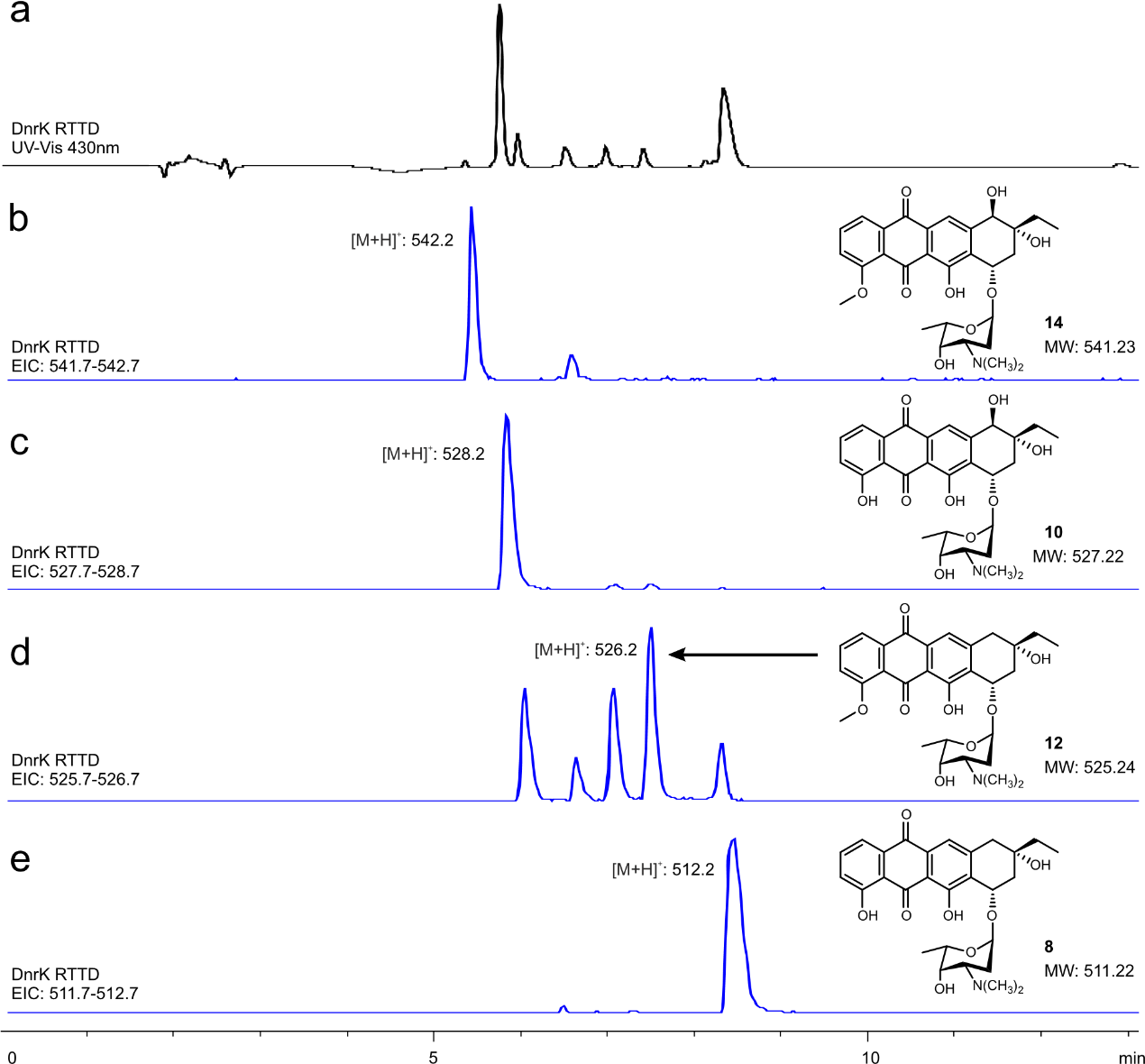

**Supplementary Figure 6. LC-MS analysis of enzymatic reactions with 5 as a substrate. a,** UV-Vis chromatogram trace recorded at 430 nm for enzymatic reaction products of DnrK RTTD. **b**, **c, d, and e,** Extracted ion chromatogram traces in positive mode for DnrK RTTD products.

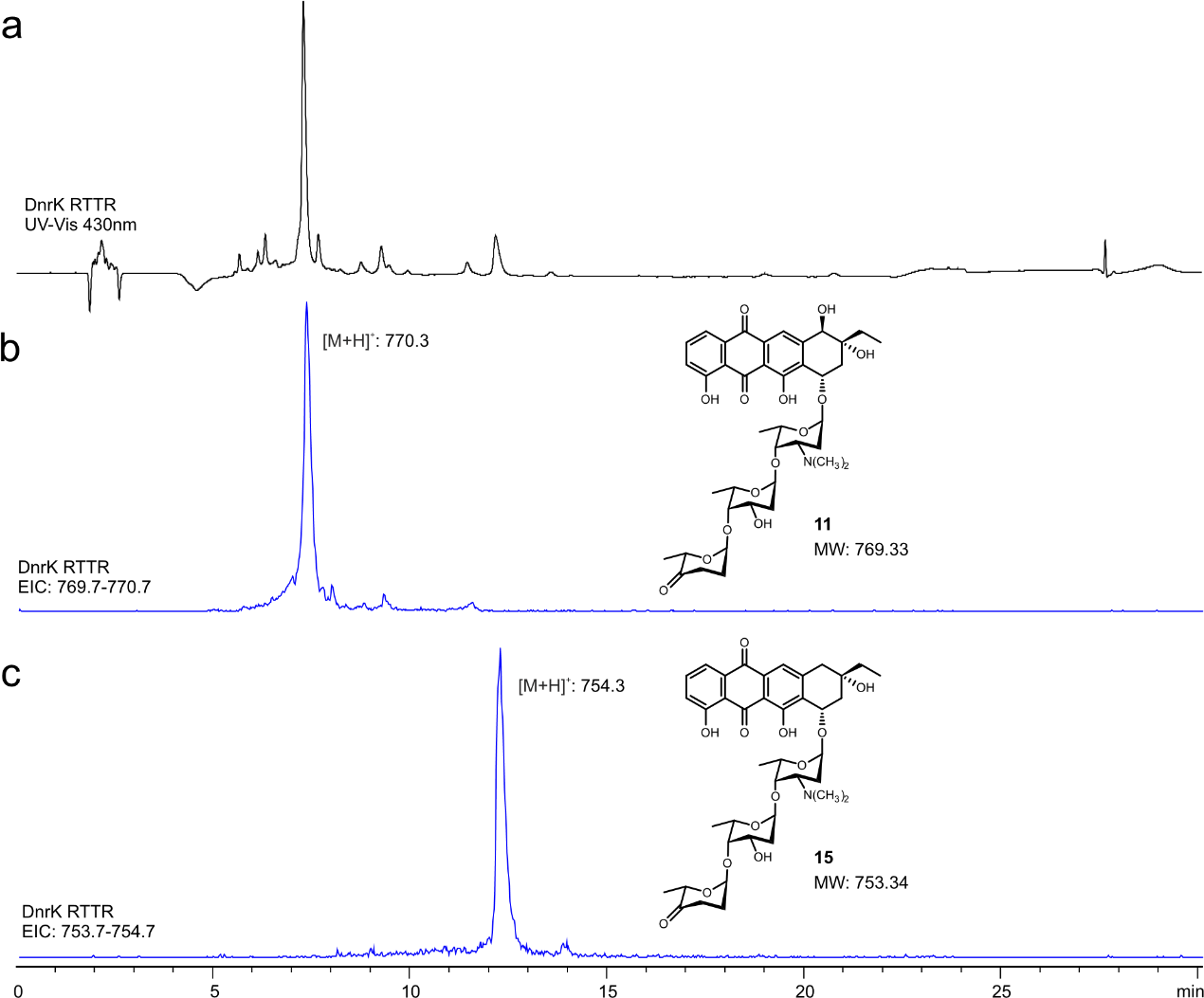

**Supplementary Figure 7. LC-MS analysis of enzymatic reactions with 6 as a substrate. a,** UV-Vis chromatogram trace recorded at 430 nm for enzymatic reaction products of DnrK RTTR. **b** **and** **c,** Extracted ion chromatogram traces in positive mode for DnrK RTTR products.

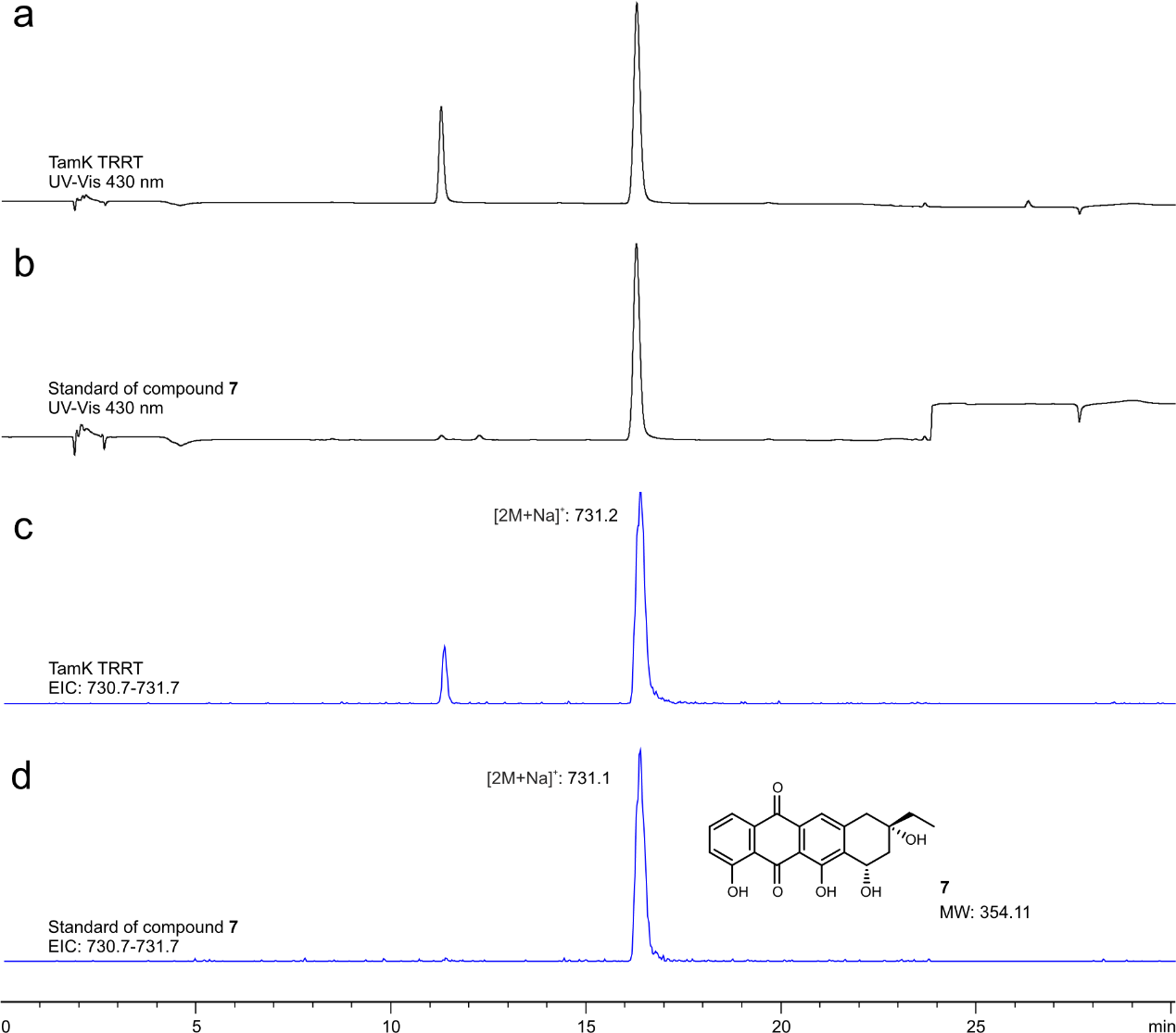

**Supplementary Figure 8. LC-MS analysis of the hydrolysed enzymatic reaction product 15 in comparison to authentic standard 7. a,** UV-Vis chromatogram trace recorded at 430 nm. Enzymatic reaction of chimeric TamK TRRT with **6** as a substrate was performed to obtain **15**, from which the glycosidic units were hydrolysed to obtain **7** (RT 16.3 min). The peak appearing at 11.3 min is a hydrolysis by-product. **b,** UV-Vis chromatogram trace recorded at 430 nm. The enzymatic reaction of native TamK with **4** as a substrate was performed to obtain the authentic standard **7**. **c,** Extracted ion chromatogram trace in positive mode of TamK TRRT products (**a**). **d,** Extracted ion chromatogram trace in positive mode of TamK product **(b)**. Products are observed as sodium adducts [2M+Na]^+^ under the conditions used, giving consistently values of 2M + 22.99.

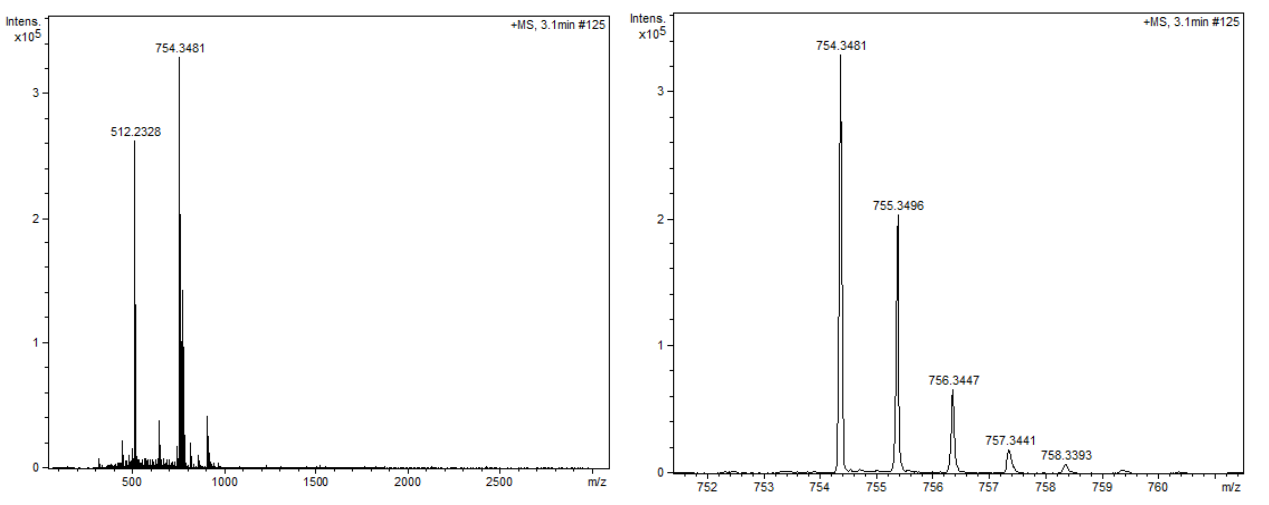

**Supplementary Figure 9. HR-MS spectrum of 15**. ESI m/z [M+H]^+^, ESI+ obs. 754.3481, calc. 754.3433.

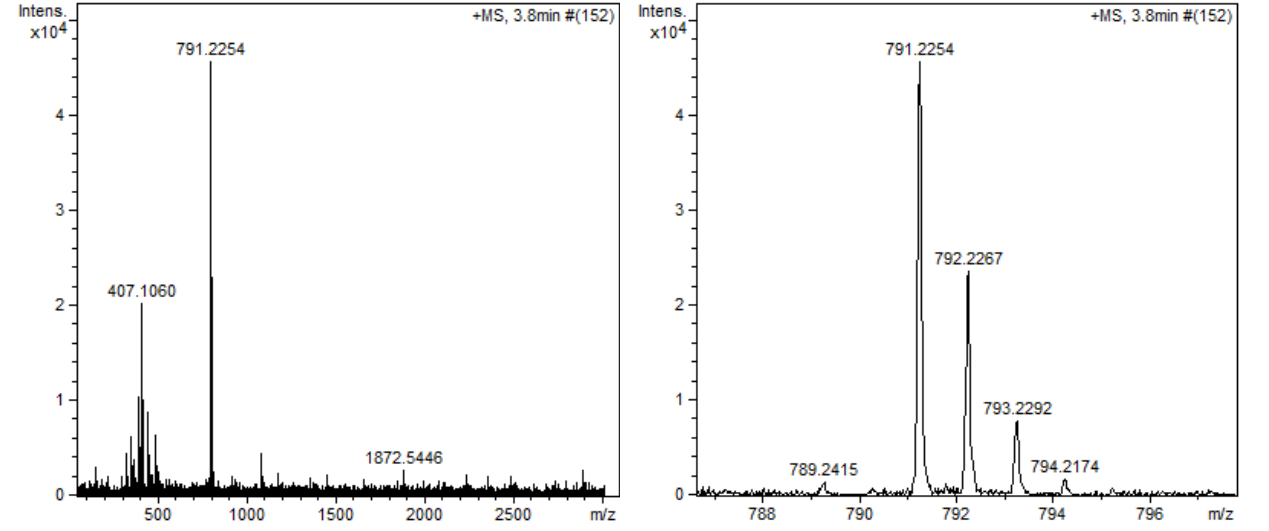

**Supplementary Figure 10. HR-MS spectrum of 13.** ESI m/z [2M+Na]^+^, ESI+ obs. 791.2254, calc. 791.2310

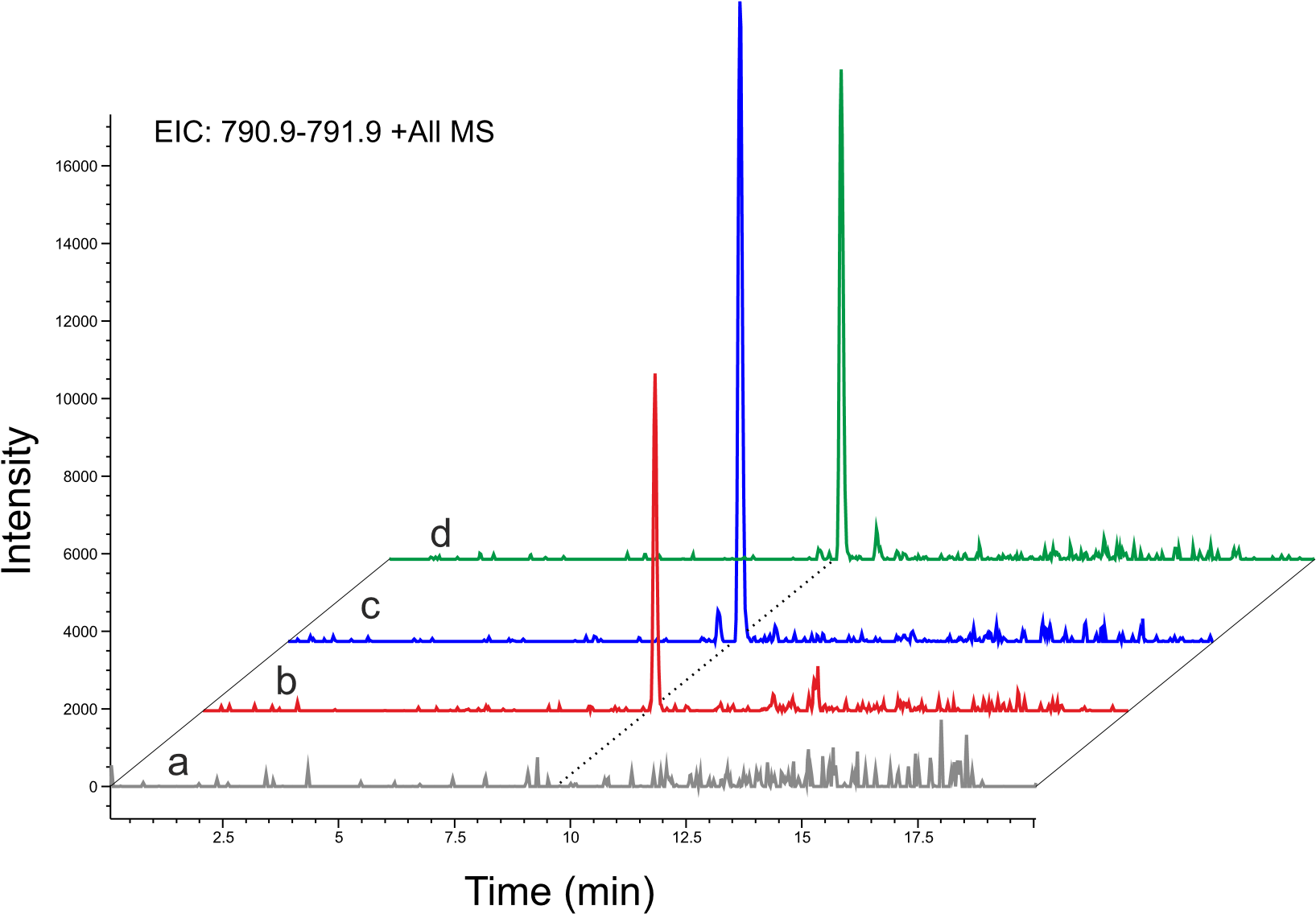

**Supplementary Figure 11. Analysis of DnrK RTTR reaction with 4 to yield 13.** **a**, Standard of compound **4.** **b,** Standard of compound **13**. DnrK and RdmB reactions were performed on **5** to obtain **14**, which was subsequently acid hydrolysed to yield the standard of compound **13.** **c**, DnrK RTTR reaction products with **4** as a substrate**.** **d,** Mixture of DnrK RTTR reaction products (**c**) and standard of compound **13** (**b**). All chromatograms are shown with the extracted ion chromatogram (EIC) for **13**. Products are observed as sodium adducts [2M+Na]^+^.

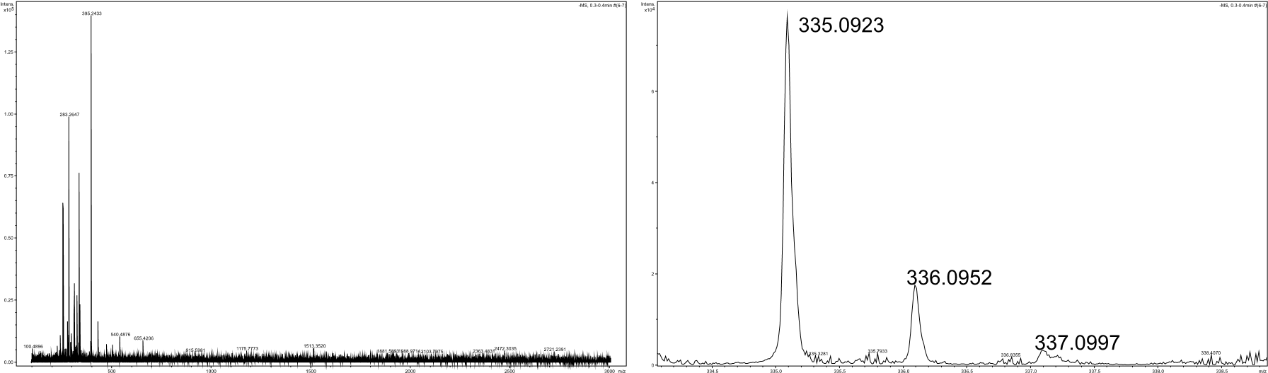

**Supplementary Figure 12. HR-MS spectrum of 16.** ESI m/z [M-H]-, ESI- obs. 335.0923, calc. 335.0925.

**Supplementary Table 2. Compound 16 recorded in CDCl_3_.** ^1^H is recorded at 500 MHz and ^13^C at 151 MHz. The 2D measurements are done either in 500 MHz or 600 MHz for ^1^H and either at 126 MHz or 151 MHz for ^13^C. The signals are internally referenced to tetramethylsilane (TMS).

| Position | δ ppm | δ ppm, *J* Hz | HMBC | COSY |
| --- | --- | --- | --- | --- |
|  | 13C | ^1^H |  |  |
| 1 | 120.0 | 7.84 dd 1.0, 7.5 | C2, C3, C4a, C12 | H2 |
| 2 | 136.9 | 7.67 dd 7.5, 8.6 | C4, C12a | H1, H3 |
| 3 | 124.7 | 7.3 dd 1.0, 8.6 | C1, C4, C4a | H2 |
| 4 | 162.5 |  |  |  |
| 4-OH |  | 12.14 s | C3, C4, C4a |  |
| 4a | 116.0 |  |  |  |
| 5 | 192.0 |  |  |  |
| 5a | 114.3 |  |  |  |
| 6 | 160.4 |  |  |  |
| 6-OH |  | 12.57 s | C5a, C6, C6a, C10a |  |
| 6a | 127.4 |  |  |  |
| 7 | 60.0 | 5.38 ddd 2.5, 5.3, 5.9 |  | H7-OH, H8 |
| 7-OH |  | 2.27 d 5.3 |  | H7 |
| 8 | 35.8 | 2.64 ddd 1.3, 5.9, 18.6 | C9, C10 | H7, H8 |
| 8 |  | 2.71 dd 2.5, 18.6 |  | H7, H8 |
| 9 | 148.6 |  |  |  |
| 10 | 119.4 | 6.44 td 1.0, 1.3 | C6a, C8, C11, C13 | H8, H13 |
| 10a | 142.5 |  |  |  |
| 11 | 118.7 | 7.61 s | C5, C5a, C6a, C10, C12 |  |
| 11a | ND |  |  |  |
| 12 | 181.9 |  |  |  |
| 12a | 133.4 |  |  |  |
| 13 | 30.6 | 2.36 2H qd 1.0, 7.4 | C8, C9, C10, C14 | H9, H14 |
| 14 | 11.5 | 1.20 3H t 7.4 | C9, C13 | H13 |

s=singlet, d=douplet, t=triplet q=quartet, ND = no data, integrals are 1 unless otherwise stated

OH

OH

O

OOH

**Supplementary Figure 13. Main HMBC and COSY correlations for 16.** HMBC correlations are presented with blue arrows and COSY correlations with pink lines.

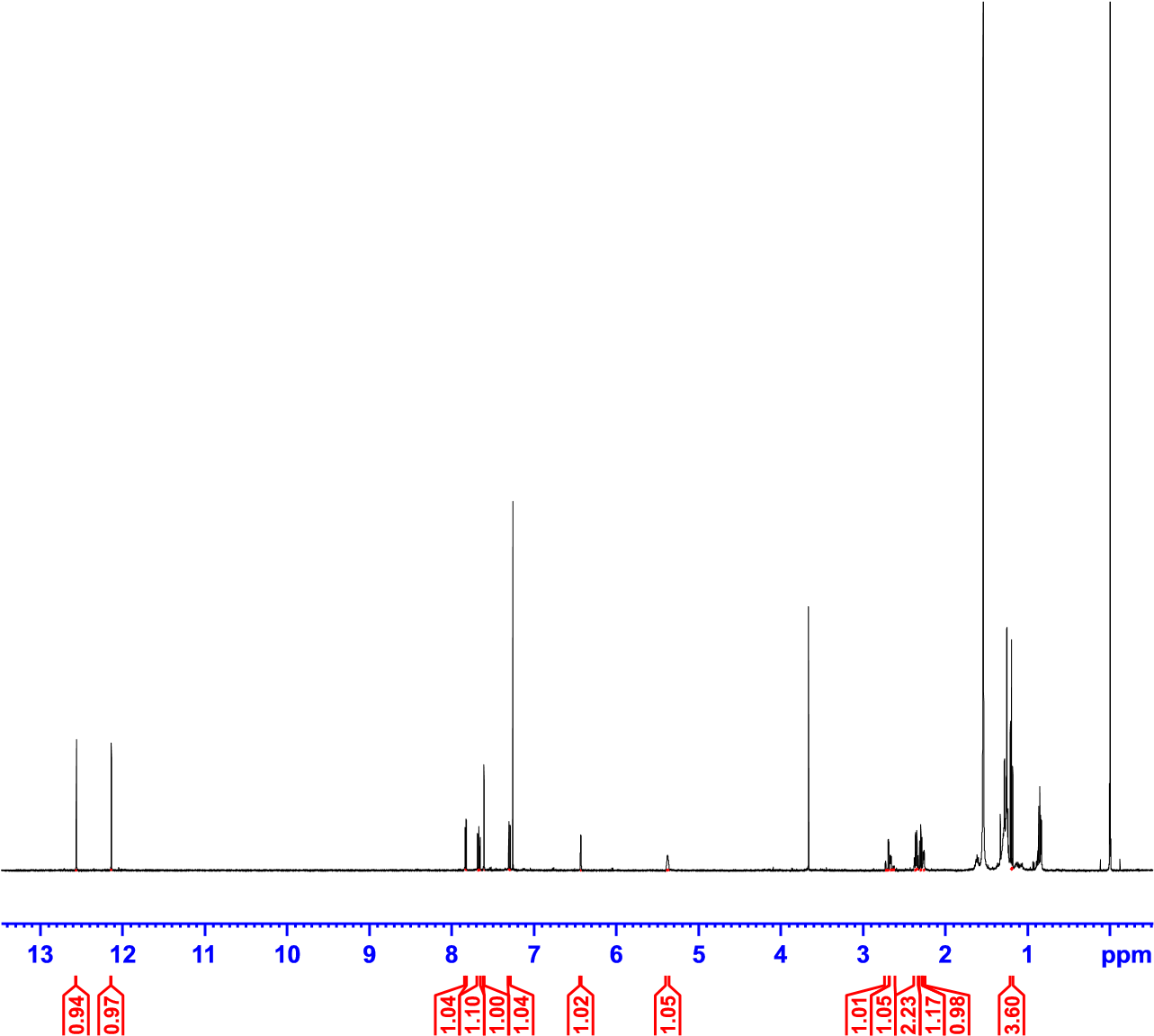

**Supplementary Figure 14. 1H spectrum of 16 in CDCl3.**

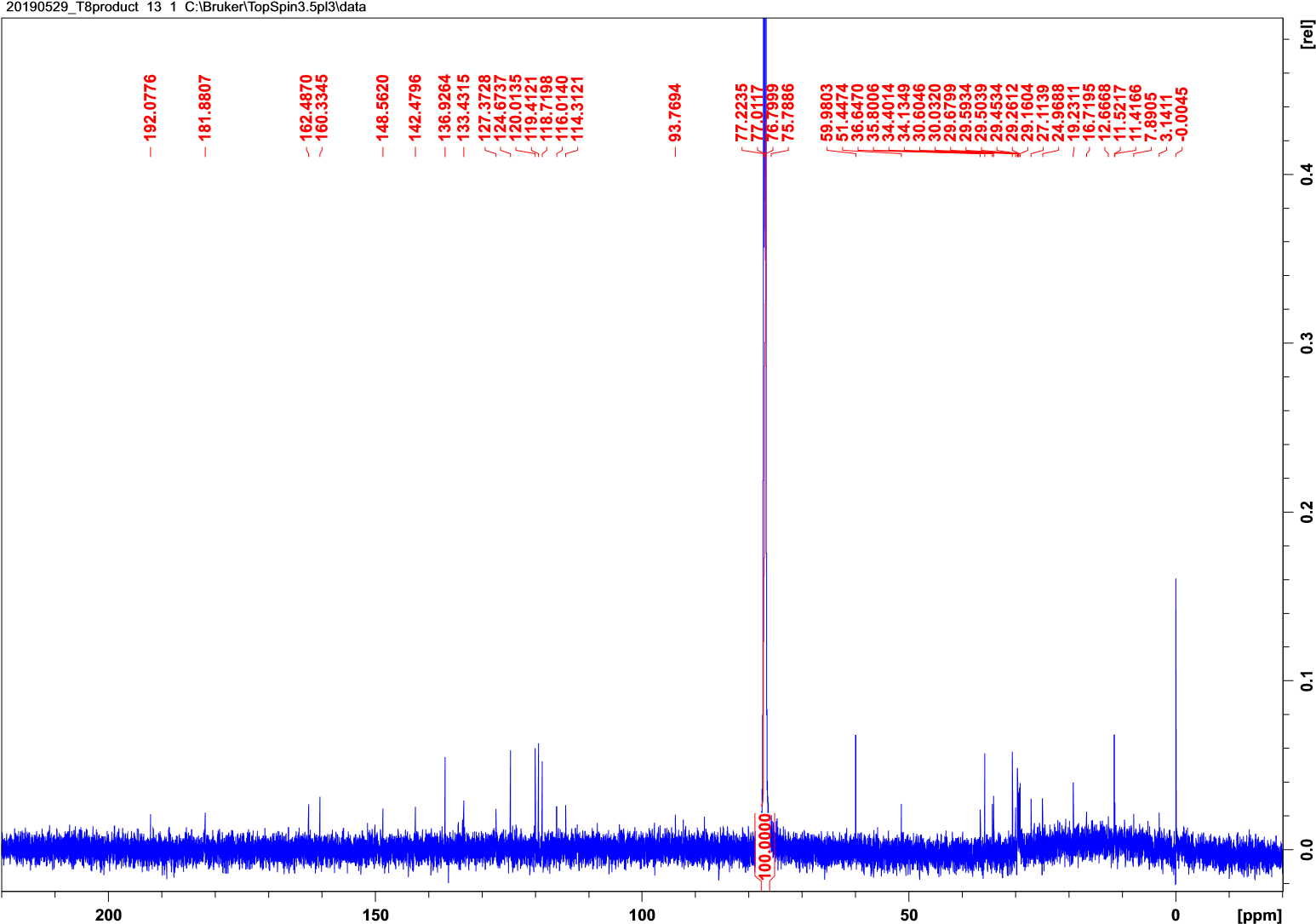

**Supplementary Figure 15. 13C spectrum of 16 in CDCl3.**

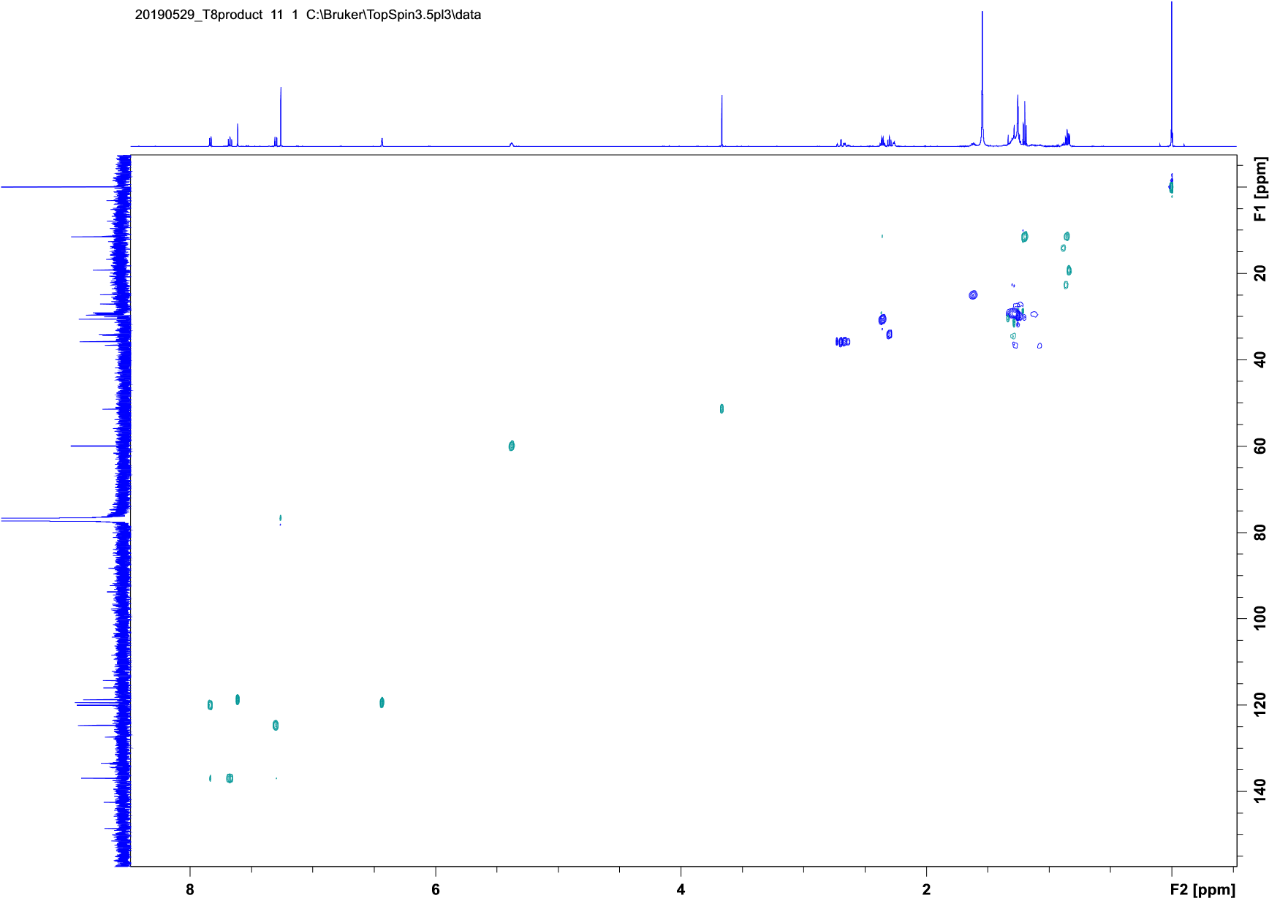

**Supplementary Figure 16. HSQCDE spectrum of 16 in CDCl3.**

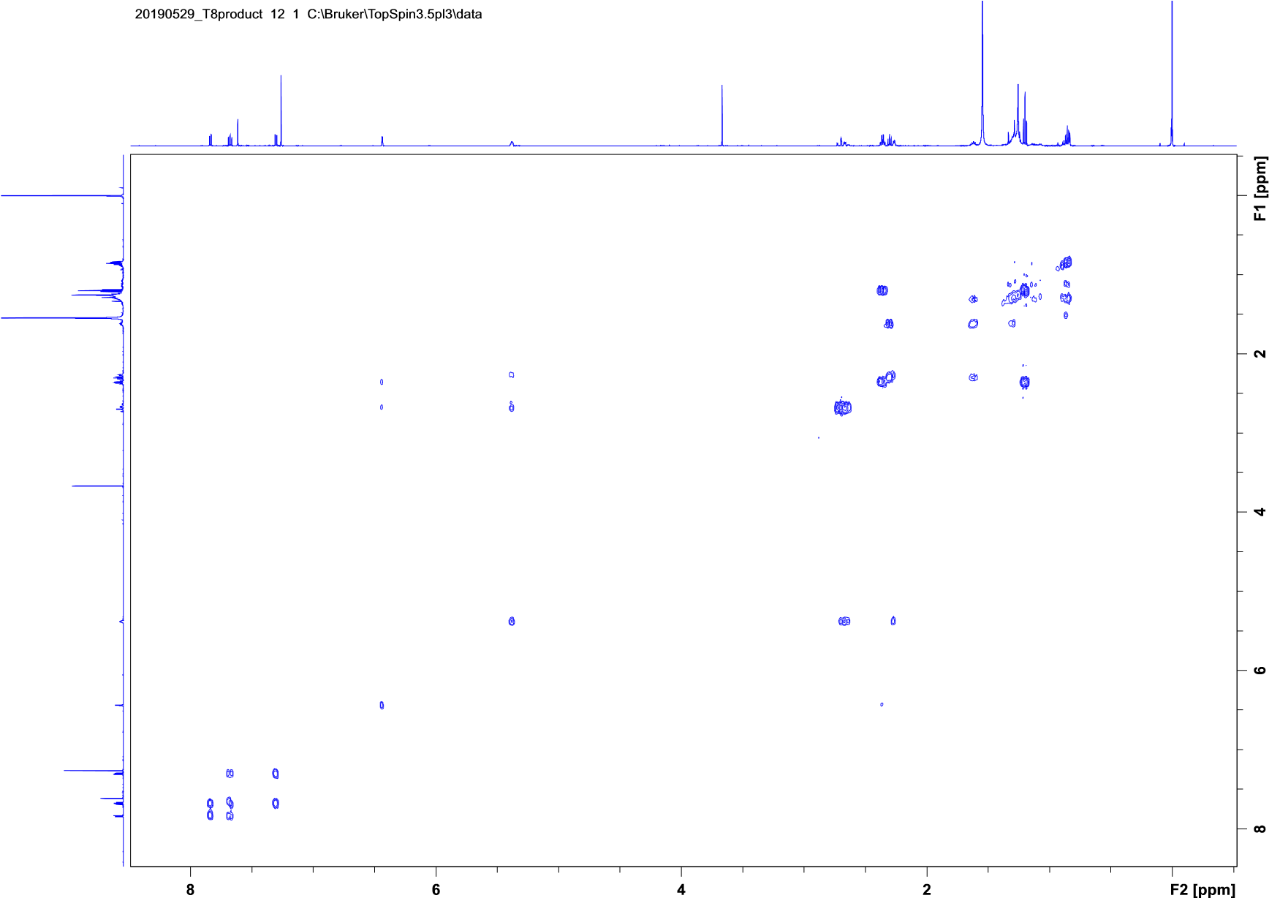

**Supplementary Figure 17. COSY spectrum of 16 in CDCl3.**

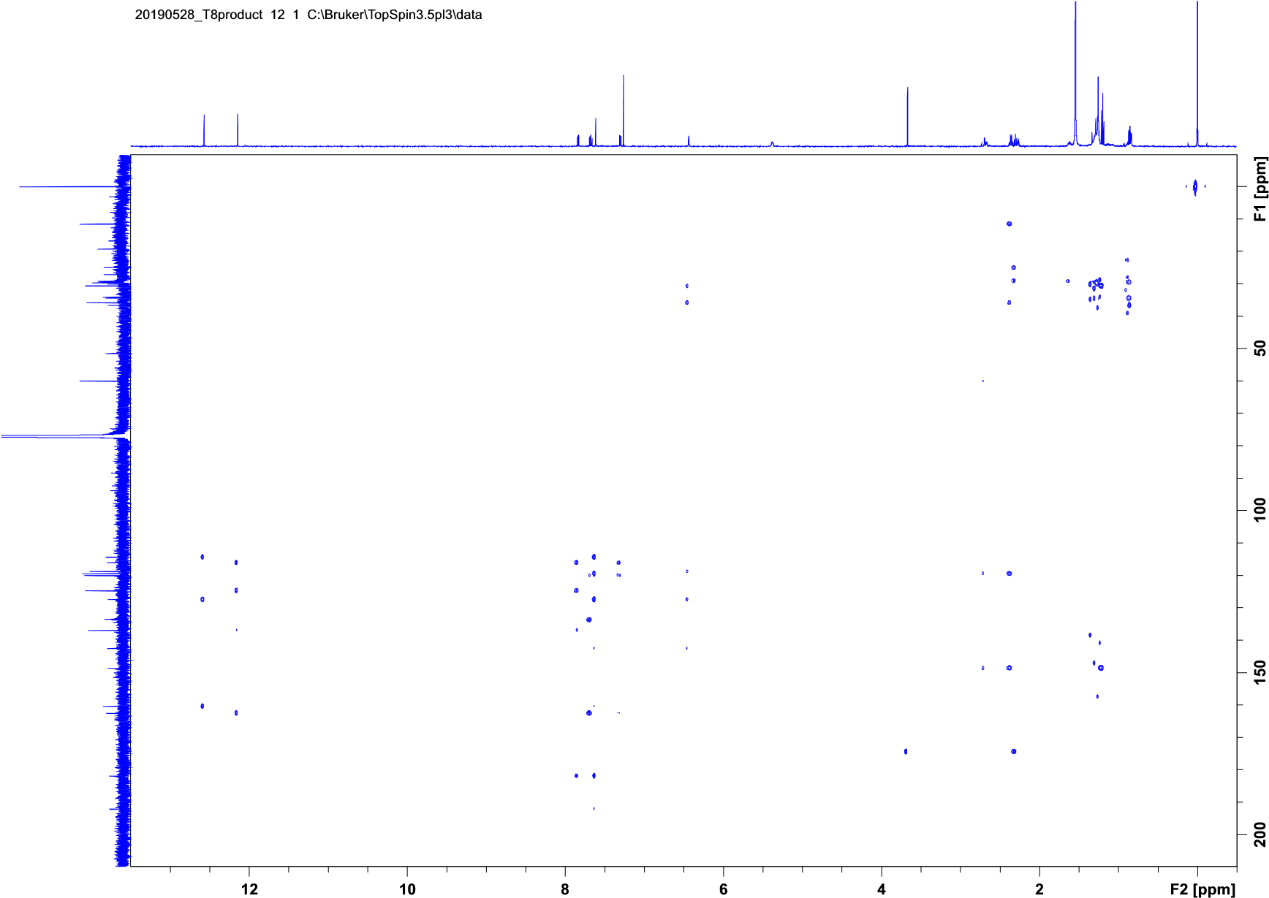

**Supplementary Figure 18. HMBC spectrum of 16 in CDCl3.**

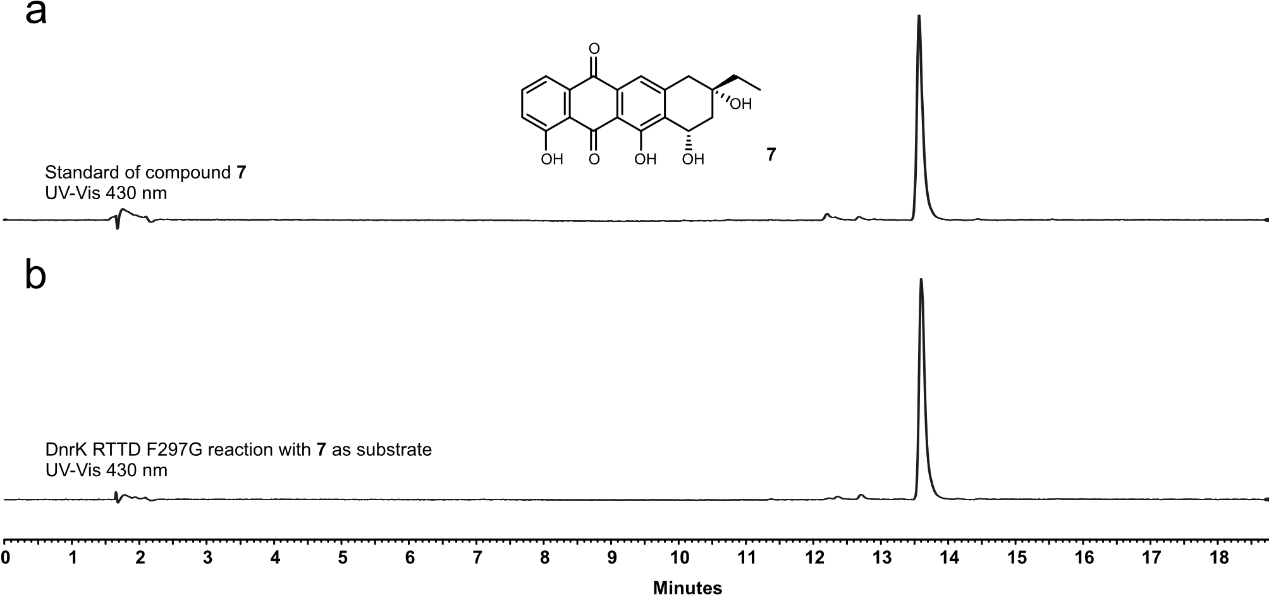

**Supplementary Figure 19. HPLC analysis of elimination reaction substrate. a,** UV-Vis chromatogram trace recorded at 430 nm for **7**. Standard of compound **7** was obtained through enzymatic reaction of TamK with **4** as a substrate. **b,** UV-Vis chromatogram trace recorded at 430 nm for DnrK RTTD F297G enzymatic reaction products with **7** as a substrate. No elimination reaction was observed.

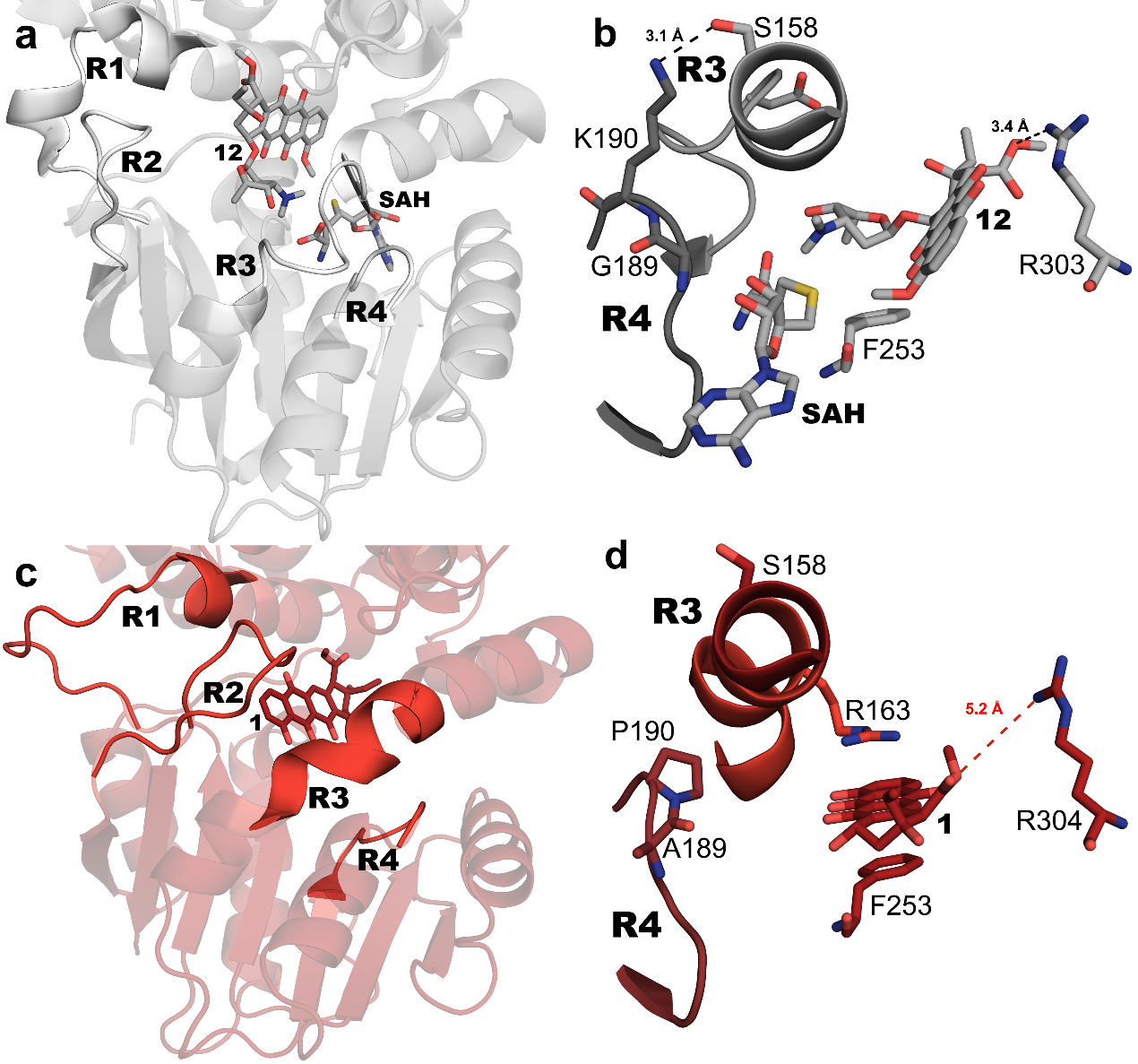

**Supplementary Figure 20.** **Structure of DnrK RTTT.** The DnrK RTTT chimera crystallized with the substrate bound between the anthracycline and SAM binding sites. **a**, Overall structure of DnrK WT**.** **b,** Interactions of regions R3 and R4 of DnrK WT**.** **c,** Overall structure of DnrK RTTT. The unique binding of the substrate occupies the SAM binding site and has led to several key changes in the structure. Due to **1** being bound further inside the active site, region R2 has shifted considerably to close the active site. In addition, the helix that forms region R3 has extended considerably**.** **d,** The interface between regions R3 and R4 is the most dissimilar in DnrK RTTT. Due to the non-reactive binding of the substrate, as indicated by the large distance to the catalytic R304, the substrate is protruding into the SAM binding site. The aglycone is placed in π-π between R163 from R3 and F253 from the main chain. The K190P mutation prevents the interaction with S158 from R3.

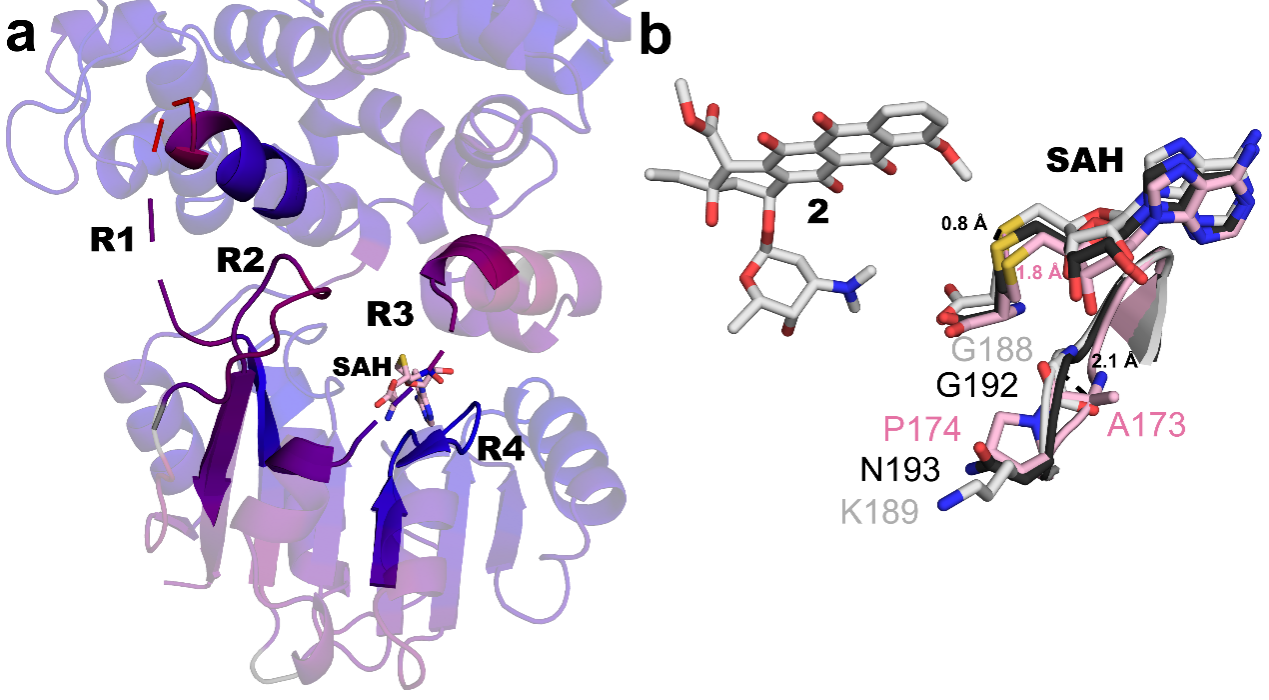

**Supplementary Figure 21. TamK structure and changes in region R4 and SAM binding. a,** The overall fold of TamK was aligned against DnrK colored by RMSD, with blue regions indicating a high degree of conservation and pink regions showing a higher variation between the two structures. The absence of substrate led to lack of density to build loops belonging to regions R1 and R3. **b,** Comparing the SAM binding region R4 of DnrK WT (white) (PDBID: 1TW2), RdmB (black) (PDBID: 1R00) and TamK WT (pink). The presence of P174 leads to a 2.1 Å rearrangement of the loop of region R4, leading to a repositioning of SAM (SAH in the figure). This movement is sufficient to prevent correct position of the cofactor in order to catalyse 4-O-methylation of the substrate.

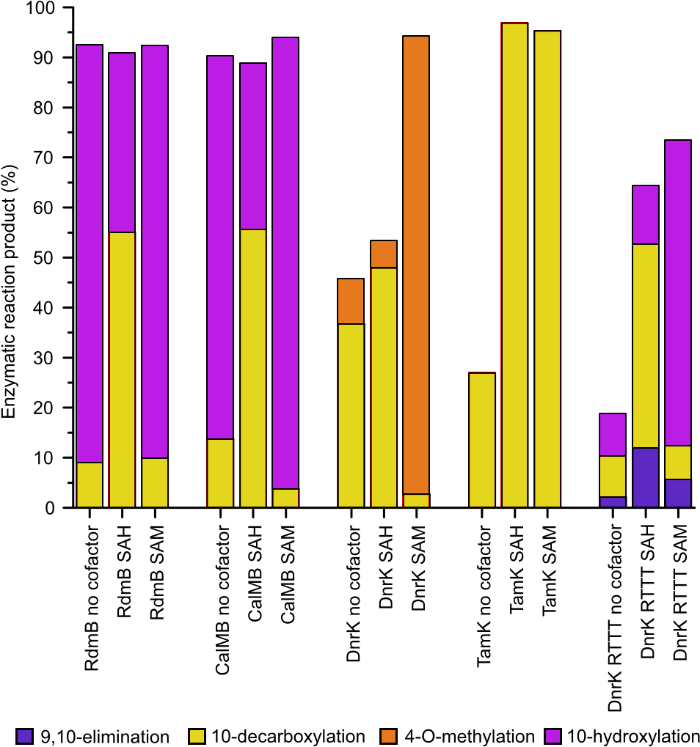

**Supplementary Figure 22. Cofactor SAM requirement enzymatic assays.** The enzymes RdmB, CalmB, DnrK, and TamK had **5** as a substrate, whereas DnrK RTTT had **4** as a substrate. The enzymatic reactions with either native or chimeric enzymes were performed with either no added cofactor, SAH added as cofactor (400 µM), or SAM added as cofactor (400 µM). The reaction product yields were calculated from HPLC chromatogram traces by normalized peaks areas. The overall percentages may not add up to 100 % in all samples meaning that some unreacted substrate remained in the reaction mixture.

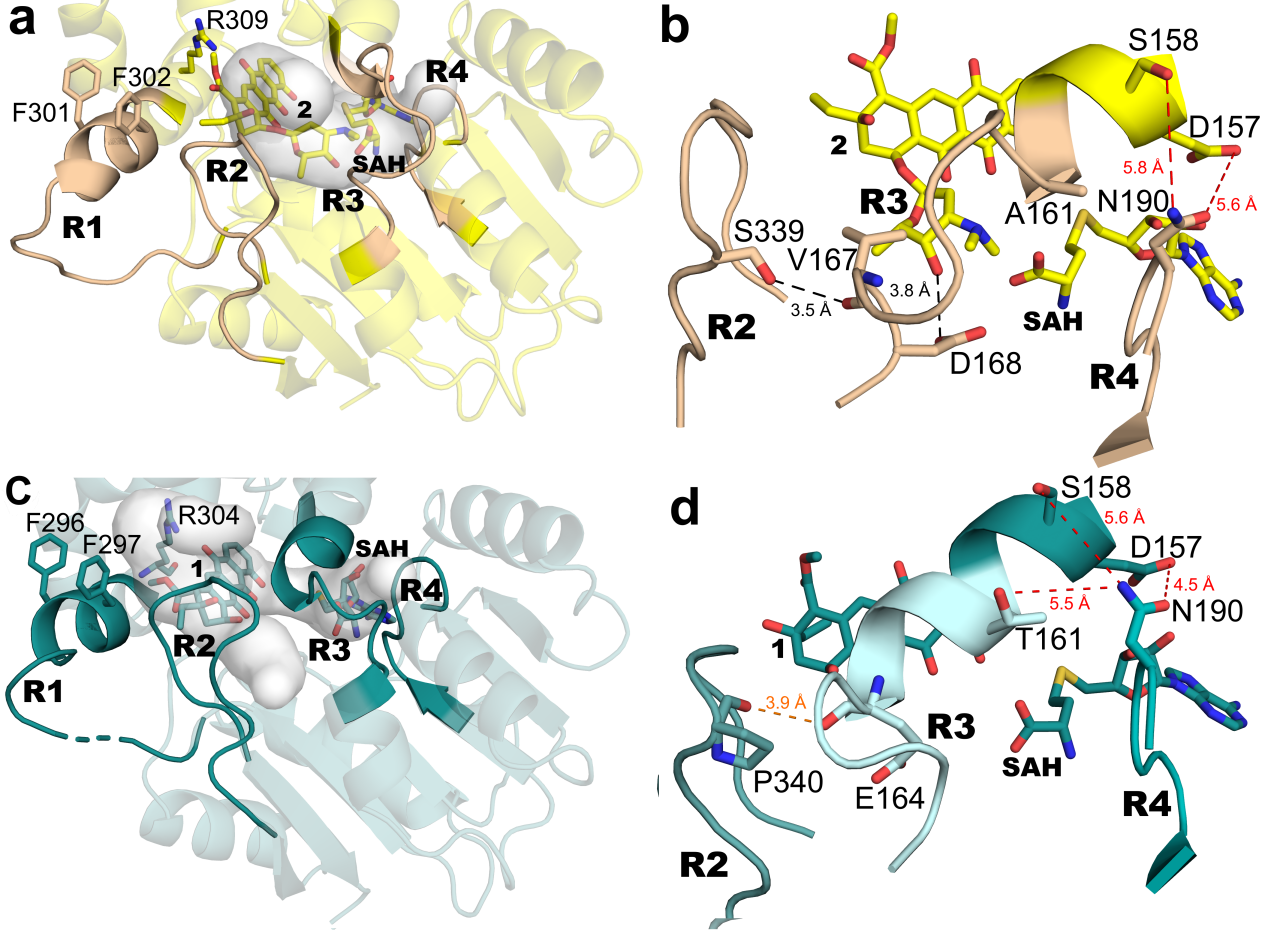

**Supplementary Figure 23.** **Comparison between DnrK RTTR and DnrK RTCR. a,** Overall fold of DnrK RTCR, crystallized in a closed conformation as evidenced by the CAVER analysis (light grey). The aromatic residues from region R1 prevent access to bulk solvent to the active site. R3 from CalMB shows the formation of a loop instead of the more common helix conformation. **b,** Closer observation of the inter-region interactions show a single hydrogen bond between S339 from R2 and the main chain of V167 in R3. D168 (R3) is seen interacting with the sugar moiety of the substrate. Similarly to DnrK RTTR (and DnrK WT) there is no hydrogen bonding possible between regions R3 and R4. **c,** Overall fold of DnrK RTTR. **d,** Inter-region interactions of DnrK RTTR.

**Supplementary Table 3. Data collection and refinement statistics**

|  | DnrK CDDD | DnrK TDDD | DnrK RTTD | DnrK RTTR | DnrK RTTD F297G | DnrK RTTT | DrnK RTCR | TamK |
| --- | --- | --- | --- | --- | --- | --- | --- | --- |
| **Data collection** |  |  |  |  |  |  |  |  |
| Space Goup | P 21 21 2 | P 1 21 1 | P 1 21 1 | P 1 21 1 | C 2 2 21 | P 1 21 1 | P 1 21 1 | C 1 2 1 |
| Cell dimensions |  |  |  |  |  |  |  |  |
| *a, b, c* (Å) | 60.80, 103.14, 66.49 | 60.5, 101.3, 62.9 | 60.47, 102.64, 121.85 | 60.01, 102.42, 122.19 | 60.81, 125.14, 102.41 | 58.80, 110.40, 64.21 | 60.40, 105.69, 64.45 | 122.84, 39.69, 97.98 |
| α, β, γ (°) | 90.0, 90.0, 90.0 | 90.0, 102.2, 90.0 | 90.0, 98.2, 90.0 | 90.0, 99.3, 90.0 | 90.0, 90.0, 90.0 | 90, 108.6, 90 | 90.0, 111.2, 90.0 | 90.0, 114.1, 90 |
| Resolution (Å) | 44.87 - 2.45 (2.51 - 2.45)) | 48.04 - 1.53 (1.59 - 1.53) | 49.80 - 2.38 (2.47 - 2.38) | 47.13 - 2.21 (2.27 - 2.21) | 34.4 - 1.68 (1.74 - 1.68) | 55.71 - 2.316 (2.399 - 2.316) | 49.7- 2.39 (2.48 - 2.39 | 28.04 - 1.51 (1.57 - 1.51) |
| R_merge_ | 13.5 (96.8) | 5.5 (58.8) | 11.6 (84.8) | 11.2 (102.7) | 7.3 (89.3) | 23.4 (100.9) | 11.6 (24.2) | 22.62 (42.5) |
| *I/*σ*I* | 9.19 (1.56) | 11.18 (1.83) | 7.4 (1.1) | 5.9 (0.9) | 13.29 (0.99) | 8.7 (2.2) | 8.3 (4.1) | 15.69 (2.24) |
| Completeness (%) | 99.12 (95.50) | 98.1 (98.0) | 98.4 (95.0) | 99.3 (97.2) | 90.41 (81.70) | 99.4 (100.0) | 99.1 (96.0) | 96.51 (85.60) |
| Redundancy | 5.69 (4.83) | 2.69 (2.56) | 4.4 (4.3) | 3.4 (3.4) | 5.5 (2.4) | 4.3 (4.4) | 3.4 (3.0) | 1.9 (1.8) |
| **Refinement** |  |  |  |  |  |  |  |  |
| No. reflections | 15808 (1483) | 109424 (10877) | 58090 (5535) | 72307 (6991) | 40509 (1086) | 33575 (3366) | 29619 (2854) | 65666 (5771) |
| *R_work_*/*R_free_* | 0.25/0.26 | 0.21/0.23 | 0.23/0.28 | 0.21/0.24 | 0.20/ 0.23 | 0.21/0.26 | 0.23/0.27 | 0.19/0.21 |
| No. atoms |  |  |  |  |  |  |  |  |
| Protein | 2521 | 5253 | 10258 | 2473 | 2578 | 5185 | 5113 | 2511 |
| Ligand/ion | 41 | 116 | 120 | 42 | 31 | 60 | 82 | 26 |
| Water | 32 | 310 | 149 | 34 | 286 | 142 | 264 | 270 |
| B-factors |  |  |  |  |  |  |  |  |
| Protein | 48.02 | 22.12 | 49.67 | 24.22 | 26.74 | 41.12 | 16.79 | 29.26 |
| Ligand/Ion | 34.98 | 33.16 | 49.62 | 48.72 | 49.13 | 20.00 | 37.39 | 24.13 |
| Water | 43.32 | 28.47 | 44.15 | 31.98 | 35.76 | 39.58 | 20.12 | 40.42 |
| R.m.s. deviations |  |  |  |  |  |  |  |  |
| Bond lengths (Å) | 0.011 | 0.006 | 0.005 | 0.006 | 0.009 | 0.012 | 0.009 | 0.007 |
| Bond angles (°) | 1.41 | 0.96 | 0.86 | 0.86 | 1.19 | 1.65 | 1.18 | 1.07 |
